# gnSPADE: a reference-free deconvolution method incorporating gene network structures in spatial transcriptomics

**DOI:** 10.1101/2025.08.26.672457

**Authors:** Aoqi Xie, Yuehua Cui

**Affiliations:** Department of Statistics and Probability, Michigan State University, East Lansing, MI

**Keywords:** Spatial deconvolution, Latent Dirichlet allocation model, Gene network, Reference-free deconvolution

## Abstract

Spatial transcriptomics (ST) technologies have offered unprecedented insights into the spatial organization of gene expression, allowing for the study of tissue architecture, domain boundaries, and cell-cell interactions. However, most ST data generated so far are at multicellular resolution, where each spot captures transcripts from a mixture of diverse cells of different cell types. While reference-based deconvolution approaches offer robust solutions, they rely heavily on the availability and quality of external single-cell reference data, which may be incomplete, unavailable, or poorly matched to the spatial data. Moreover, even when such references are available, they often represent only broad cell types, potentially obscuring finer subpopulation structures and masking intra-type heterogeneity. To overcome these limitations, we introduce gnSPADE, a reference-free spatial deconvolution method that incorporates gene network structures via a Markov random field within a latent Dirichlet allocation (LDA) modeling framework. gnSPADE jointly infers cell type-specific transcriptional profiles and spatial compositions without external references. Applied to synthetic and real ST datasets, gnSPADE achieves improved accuracy, spatial resolution, and biological interpretability compared to other methods, highlighting the power of reference-free deconvolution in resolving complex tissues.

## 1 Introduction

Spatial transcriptomics (ST) has emerged as a transformative approach for mapping gene expression within the spatial context of intact tissue, offering critical insights into tissue architecture, cellular organization, and disease pathology[1, 2, 3]. By capturing spatially resolved transcriptomic data, ST enables the discovery of spatially variable genes (SVGs) and the identification of tissue-specific transcriptional patterns that are often obscured in dissociated single-cell datasets[4, 5]. Recent advances such as Xenium and other high-resolution imaging-based platforms[6, 7] have pushed the boundary toward single-cell spatial transcriptomics; however, their gene coverage remains limited, typically profiling a few hundred genes, compared to the *>*10,000 genes accessible through other ST platforms[8, 9]. This reduced transcriptome depth constrains the detection of novel marker genes and restricts the scope of downstream analyses such as cell state characterization and trajectory inference. Furthermore, identifying cell-type-specific SVGs, as implemented in tools like STANCE[10], necessitates an accurate cell type deconvolution step, highlighting the foundational role of deconvolution in ST workflows. Robust deconvolution remains critical not only for enhancing interpretability but also for enabling cell-type-resolved functional analyses and disease-associated spatial mapping.

Reference-based deconvolution methods, such as RCTD[11], Cell2location[12], SPOTlight[13], and CARD[14], have proven effective for spatial transcriptomics by leveraging annotated single-cell RNA sequencing (scRNA-seq) datasets to infer cell-type composition in spatial data[15, 16]. However, their reliance on high-quality external reference datasets poses a critical limitation. In many cases, appropriate reference datasets are either unavailable or poorly matched, leading to biased or noisy estimates, particularly when batch effects or platform discrepancies distort cell-type signatures[17, 18]. Moreover, reference datasets often capture only a limited repertoire of major cell types, lacking resolution at the level of finer-grained sub-cell populations[19, 20]. This limitation is especially problematic in heterogeneous tissues, where biologically meaningful subtypes, despite being functionally distinct, may escape detection due to insufficient marker gene annotation or low representation in scRNA-seq atlases. Therefore, reference-free deconvolution strategies are increasingly necessary to resolve latent cellular subtypes directly from spatial transcriptomics data. Such approaches enable data-driven discovery of novel cell states and sub-cell types, even in the absence of prior knowledge.

With the growing recognition of limitations in reference-based approaches, several reference-free deconvolution methods have been proposed to analyze ST data in an unsupervised manner. Among them, SpiceMix[21] and STdeconvolve[22] represent two widely used models. SpiceMix employs a non-negative matrix factorization (NMF) framework augmented with spatial priors encoded via hidden Markov random fields (HMRF), enabling spatial smoothing of inferred components[21]. However, it is not specifically optimized for spatial cell-type deconvolution and lacks an intrinsic criterion for selecting the optimal number of latent cell types, which can limit biological interpretability. In contrast, STdeconvolve is grounded in latent Dirichlet allocation (LDA) model[23], a probabilistic topic modeling approach that decomposes the transcriptomic profiles into latent topics, interpreted as cell types, and estimates both the cell type-specific gene expression signatures and their spatial compositions[22]. Despite its utility, STdeconvolve inherits a core limitation of classical LDA: the assumption of gene independence. This simplification ignores gene-gene interactions and co-expression patterns, which are fundamental to cellular function[24]. In reality, gene expression is coordinated through regulatory networks, and sets of co-expressed genes often define cell types more robustly than single-gene signatures[25, 26]. These observations underscore the need for deconvolution models that explicitly incorporate gene-gene relationships, motivating the integration of gene connectivity information into future deconvolution frameworks.

Here, we present **gnSPADE**, a method that incorporates **g**ene **n**etwork structures into **SPA**tial **DE**convolution of ST data. It is an extension of the LDA framework that incorporates gene-gene interaction networks for ST deconvolution. gnSPADE enhances the generative topic modeling process by imposing a Markov random field (MRF) on the latent gene-cell type layer[27], enabling the model to integrate structured gene information, such as protein-protein interaction (PPI) networks[28], gene regulatory networks (GRNs)[29], and other biological pathways[30, 31]. By encouraging genes connected within these structures to share the same latent topic, gnSPADE promotes the assignment of functionally related genes to the same cell type, thereby generating more coherent and biologically meaningful cell type definitions than methods assuming gene independence. We comprehensively evaluated gnSPADE across both simulated and real datasets. Specifically, its advantages were demonstrated using two model-based simulation datasets and two single-cell-derived synthetic spatial transcriptomics datasets. Moreover, gnSPADE’s robustness was further validated through superior performance across four publicly available spatial transcriptomics datasets, spanning diverse technologies, spatial resolutions, and tissue architectures.

## 2 Methods

### 2.1 Overview of gnSPADE

#### 2.1.1 gnSPADE modeling

gnSPADE extends the standard LDA model by incorporating a MRF[27] over the latent cell type assignments, thereby integrating gene-gene correlation information to better resolve latent cell types within multicellular ST data. In this context, a tissue slice is partitioned into *D* spatial locations (i.e., spots), where each spot *d* ∈ {1, …, *D*} contains a mixture of cells from *K* distinct cell types. Each observed transcript, indexed by its unique molecular identifier (UMI), is associated with a gene *w*_*d,n*_ ∈ {1, *· · ·*, *V*}, where *V* is the total number of genes, and is assumed to originate from a latent cell type *z*_*d,n*_ ∈ {1, …, *K*}. Under this generative model, the cell type assignment *z*_*d,n*_ for each transcript in spot *d* is drawn from a multinomial distribution parameterized by *θ*_*d*_, the cell type composition in that spot. Conditional on the cell type *k*, the observed gene *w*_*d,n*_ is drawn from a multinomial distribution over genes with parameter *β*_*k*_, representing the gene expression profile of cell type *k*. The overall model framework is summarized in Fig.1. The key components of the model are defined as follows:

- ***θ*** = (*θ*_1_, *· · ·*, *θ*_*D*_)^*T*^: a *D × K* matrix, where each row *θ*_*d*_ ∼ Dir(*α*), represents the proportion of cell types within spot *d*, with 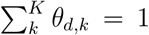. *α* is a uniform scaling parameter for the Dirichlet prior.
- ***β*** = (*β*_1_, *· · ·*, *β*_*K*_)^*T*^: a *K × V* matrix, where each row *β*_*k*_ ∼ Dir(*η*), represents the gene expression frequencies for cell type *k*, with 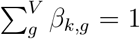. *η* is a uniform scaling parameter for the Dirichlet prior.
- *z*_*d,n*_ ∈ *{*1, *· · ·*, *K}*: the latent cell type assignment for the *n*th observed UMI in spot *d*, drawn as *z*_*d,n*_ ∼ Multinom(*θ*_*d*_), where *n*∈ *{*1, …, *N*_*d*_*}* and *N*_*d*_ is the total UMI counts in spot *d*.
- *w*_*d,n*_ ∈ *{*1, *· · ·*, *V}*: the observed gene for the *n*th UMI in spot *d*, drawn from Multinom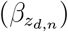, conditional on the inferred cell type.

**Figure 1:**
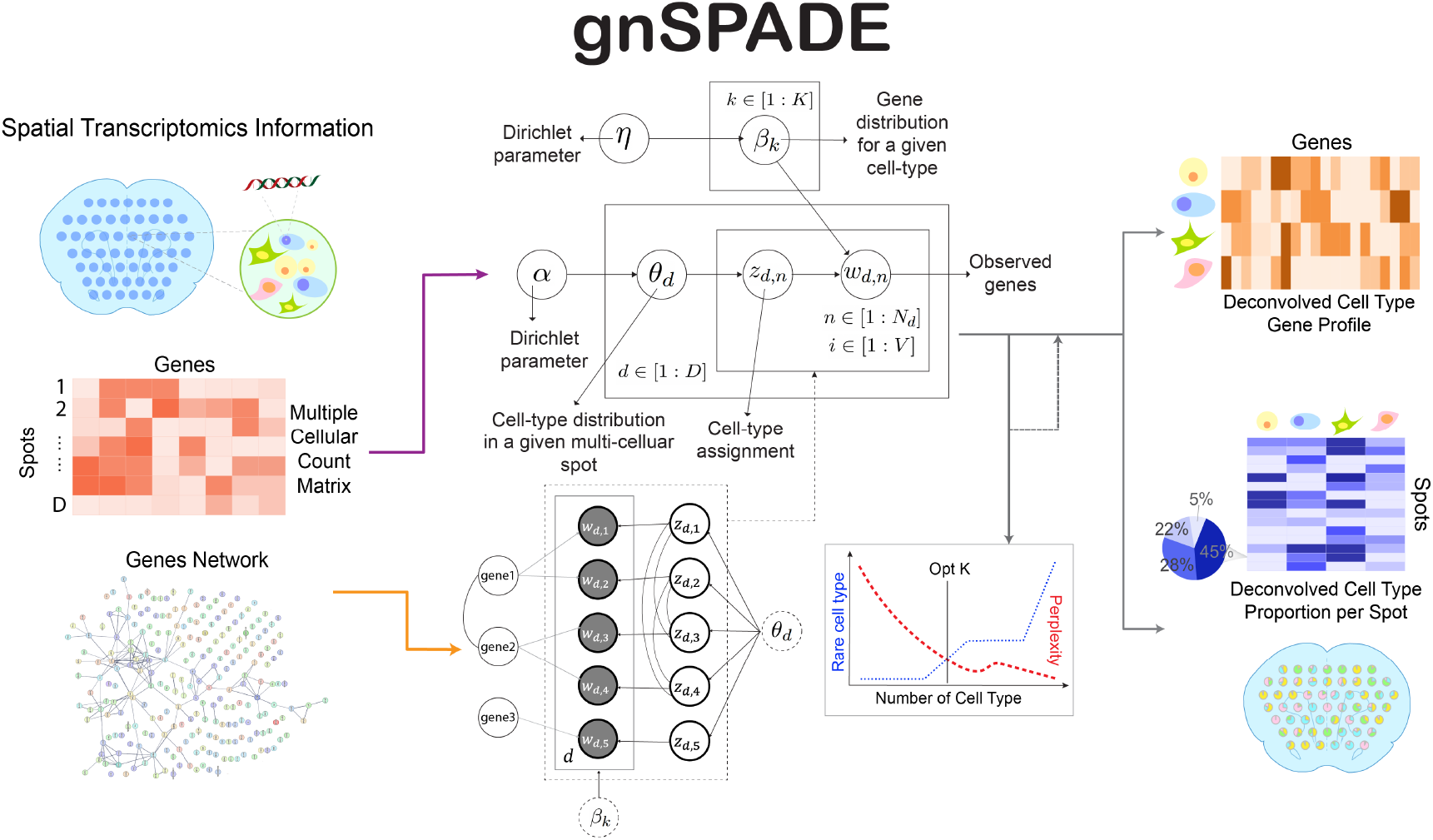
Overview of gnSPADE. gnSPADE takes a spatial transcriptomics count matrix alongside a gene-gene network graph as input. In the count matrix, rows represent spatial spots and columns represent gene expression levels. The accompanying gene network is modeled as a graph where nodes denote individual genes, and edges indicate correlation or known interactions, reflecting functional relationships such as co-expression, regulatory links, or shared pathway membership. These two data modalities are integrated within an unsupervised topic modeling framework to jointly infer two key outputs: (1) a cell type-by-gene matrix representing the transcriptional profiles of latent cell types, and (2) a spot-by-cell type matrix denoting the spatial distribution of these inferred cell types. A comprehensive formulation of model components, assumptions, and inference procedures is detailed in the Methods section.

To infer the latent parameters ***θ*** and **z**, we compute the posterior distribution conditioned on the observed gene expression data.

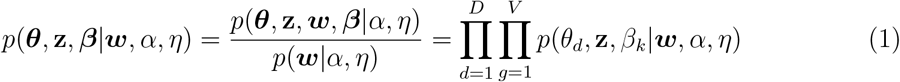

Specifically, we adopt a collapsed Gibbs sampling strategy[32][33], a standard inference method for LDA, in which the multinomial parameters ***θ*** and ***β*** are analytically integrated out. This yields a posterior distribution over the latent cell type assignments *z*_*d,n*_ that depends only on the current assignments of all other variables. Under default hyperparameter settings of *η* = 0.01 and *α* = 1*/K*, the conditional posterior probability for the latent variable *z*_*d,n*_ is given by:

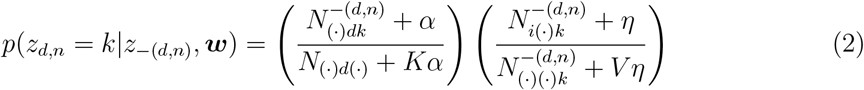

where,

- *N*_*gdk*_: the number of times gene *g* in spot *d* is assigned to cell type *k*.
- 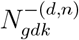: the same count excluding the current observation *w*_*d,n*_.

During each iteration of Gibbs sampling, a new assignment for *z*_*d,n*_ is sampled for each UMI *w*_*d,n*_, based on the current state of all other assignments. After a sufficient number of burn-in iterations and convergence, we estimate the cell type composition 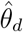 and the gene expression profile 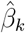 from the final latent assignments as:

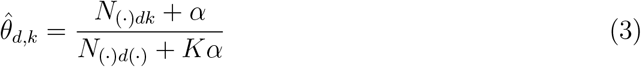

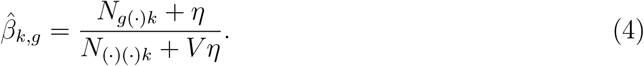

In many biological contexts, prior knowledge about gene-gene correlations is readily available and can provide valuable information for improving cell type deconvolution. For example, *Plp1* and *Mbp* are two key genes involved in the formation and maintenance of myelin sheaths in the central nervous system (CNS), and are known to be highly co-expressed in oligodendrocytes, the myelinating cells of the CNS[34, 35]. Incorporating such known co-expression patterns into deconvolution models can enhance the coherence of inferred cell types and facilitate more accurate downstream cell type annotation and biological interpretation.

To integrate gene-gene correlation information, gnSPADE defines a Markov random field (MRF)[27, 36] over the latent cell type layer. Given a spot *d* with *N*_*d*_ UMI counts, we examine all transcript pairs (*w*_*d,i*_, *w*_*d,j*_). If the two transcripts from different genes are known to be correlated based on prior knowledge (e.g., co-expression or functional interaction), we introduce an undirected edge between their respective latent cell type assignments (*z*_*d,i*_, *z*_*d,j*_). This results in an undirected graph *𝒢*_*d*_, where the nodes correspond to the latent variables 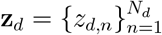, and edges connect pairs of assignments associated with correlated genes. For instance, in the example illustrated in Fig.1, the graph *𝒢*_*d*_ includes five nodes *z*_*d*,1_, *z*_*d*,2_, *z*_*d*,3_, *z*_*d*,4_, *z*_*d*,5_ and four edges linking correlated gene-cell type assignment pairs (*z*_*d*,1_, *z*_*d*,3_), (*z*_*d*,1_, *z*_*d*,4_), (*z*_*d*,2_, *z*_*d*,3_), (*z*_*d*,2_, *z*_*d*,4_). To convert the graph 𝒢_*d*_ into an MRF, we define binary edge potentials that favor similar assignments for correlated genes. Specifically, the potential between a connected pair (*i, j*) ∈ *𝒫*_*d*_ is given by:

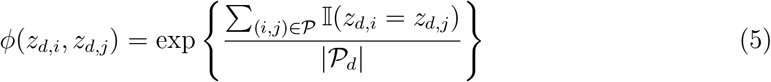

Here, *𝒫*_*d*_ denotes the set of all edges in the gene correlation graph *𝒢*_*d*_ for spot *d*, and |*𝒫*_*d*_| indicates the total number of such edges. The function 𝕀 (*·*) is the indicator function, which encourages correlated genes to be assigned to the same latent cell type. Under this formulation, the joint probability of the latent cell type assignments within spot *d* becomes:

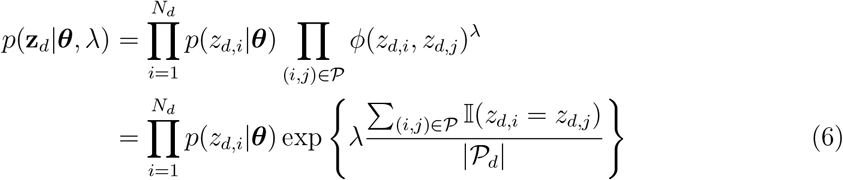

where *λ* ≥ 0 is a regularization parameter that controls the strength of the MRF prior. Then we have:

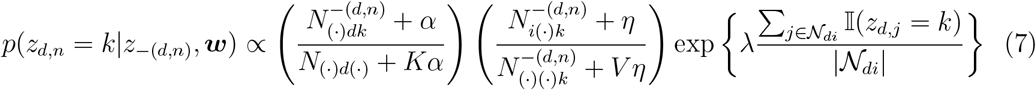

where *𝒩*_*di*_ denotes the genes that are labeled to be similar to gene *i* in the *d*th spot, |*𝒩*_*di*_| is the number of transcript counts in *𝒩*_*di*_. When *λ* = 0, gnSPADE reduces to the standard LDA model in equation (2), in which each *z*_*d,n*_ is conditionally independent given *θ*_*d*_. In contrast, gnSPADE explicitly encourages the assignments of correlated genes to align, introducing structured dependencies that reflect biological priors. Additional technical details can be found in the online supplementary file.

#### 2.1.2 Gene selection

Regardless of whether a deconvolution method is reference-based or reference-free, cell type-specific marker genes consistently provide essential biological signals that improve the accuracy and interpretability of deconvolution results[37]. To enhance computational efficiency and reduce noise from ubiquitously expressed genes (e.g., housekeeping genes), it is critical to prioritize genes that exhibit cell-type-specific expression patterns and contain meaningful biological variation.

Our preprocessing pipeline begins by removing genes with low detection rates, specifically those expressed in fewer than 1% of spatial spots. We further assume that cell-type-specific expression will manifest as overdispersed gene expression across spots, reflecting the heterogeneous spatial distribution of distinct cell types. Therefore, we retain genes with significantly overdispersed genes[38] with higher-than-expected variance, indicative of biologically meaningful transcriptional heterogeneity. Including too many genes in the input matrix can hinder convergence of topic models and reduce interpretability. To ensure model tractability and prevent overfitting, we limit the input to the top 1,000 most overdispersed genes, which are expected to capture the majority of cell-type-defining signals, especially considering that the number of marker genes relevant to most biological questions typically falls well below this threshold. Users may also supplement the selected genes with additional cell-type-specific markers or genes of interest based on prior biological knowledge or biological hypotheses, allowing for customized refinement of the deconvolution process.

#### 2.1.3 Gene-gene network

Recent years have seen the rapid accumulation of high-confidence biological networks, including protein-protein interaction (PPI) networks, metabolic pathways, and gene regulatory networks. These curated resources have been widely leveraged to enhance gene expression analyses, particularly in bulk and single-cell transcriptomics[39, 40]. However, the integration of such network information into spatial transcriptomics deconvolution remains underexplored.

In this study, we incorporate the PPI networks extracted from the STRING database[28], to guide the modeling of gene-gene dependencies. PPI networks have been extensively used in cancer research, particularly in pancreatic cancer, to uncover mechanistic insights into genomic alterations and cellular organization[41]. For network construction, STRING serves as our primary source, while additional resources such as the Kyoto Encyclopedia of Genes and Genomes (KEGG)[30] offer complementary views of functional gene relationships. The biological relevance of our network-informed deconvolution is further supported through literature curation, Reactome[31], Gene Ontology (GO) and other enrichment analysis[42].

To enhance model flexibility, users may also supplement the gene interaction network with prior biological knowledge or hypothesis-driven gene sets, enabling customized refinement of the deconvolution process.

#### 2.1.4 Selecting the number of cell types

In standard LDA modeling, perplexity is a widely used metric to assess model fit, with lower perplexity indicating better predictive performance[23]. Similar to STdeconvolve[22], we determine the optimal number of deconvolved cell types by jointly considering perplexity and the number of rare cell types, defined as those with an average proportion of less than 5% across spatial spots. The perplexity measure across *D* spots is defined as:

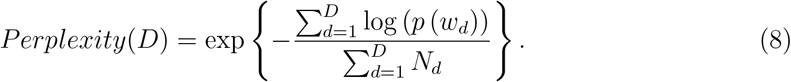

To identify the optimal number of cell types *K*, we perform a grid search starting from *K* = 2, incrementally increasing *K* based on the complexity of the tissue and any available prior biological knowledge. While perplexity typically decreases with increasing *K*, overfitting becomes a concern as the model begins to capture noise rather than signal. To mitigate this, we monitor the emergence of rare cell types, which tend to increase in number with larger *K* and are often associated with reduced deconvolution accuracy when their mean abundance falls below 5%. This 5% threshold reflects empirical observations but can be adjusted depending on the analytical goal, particularly when identifying biologically meaningful rare cell types. We select the optimal *K* by balancing these two criteria, minimizing perplexity while avoiding excessive fragmentation into rare, low-confidence cell types.

### 2.2 Evaluation Metric

#### 2.2.1 Annotation of deconvolved cell types

In the model-free ST simulation, we generated synthetic spot-level data derived from a single-cell RNA-seq dataset, for which the true cell type composition per spot is known. To approximate the ground truth transcriptional profiles of each cell type, we computed the average expression of each gene across all cells belonging to that cell type. To assess the performance of gnSPADE and other deconvolution methods, we evaluated how well the inferred cell types match the true ones by calculating the maximum Pearson correlation between each deconvolved transcriptional profile and the ground truth profiles.

When applying gnSPADE to real ST datasets, interpreting the identity of deconvolved cell types is essential. Our post hoc interpretation strategy involves two key steps:

- **Gene filtering and cell type interpretation:** To identify genes most representative of each deconvolved cell type, we first focus on those with consistently high expression within each inferred profile. After selecting the optimal number of cell types, we normalize the gene expression counts within each deconvolved cell type such that the total expression sums to 1,000, approximating a pseudo-cellular transcriptome. Genes with normalized counts of at least 5 are retained as high-frequency genes and used to support the interpretation of each deconvolved cell type, ensuring that only biologically meaningful contributors are considered.
- **Log2 fold-change analysis for sub-cell type identification:** To refine the characterization of deconvolved cell types and identify potential subtypes, we compute the log2 fold-change of each gene’s expression in a given cell type relative to its mean expression across all other cell types. Genes with log2 fold-change *>*1 are considered potentially differentially expressed and are used to define the unique transcriptional signature of that cell type. These marker genes are then subjected to Over-Representation Analysis (ORA) to evaluate enrichment in known biological processes or pathways, thereby aiding the functional annotation of each deconvolved population[42][43].

#### 2.2.2 Comparison of different methods

To systematically compare the performance of different deconvolution methods, particularly on datasets with ground truth, we evaluate two key aspects: (1) the accuracy of deconvolved cell type transcriptional profiles, and (2) the accuracy of cell type compositions across spatial spots.

For transcriptional profiles, we compute both Pearson and Spearman correlation coefficients between each ground truth cell type and its best-matched deconvolved counterpart. Higher correlation values indicate greater similarity to the ground truth, reflecting more accurate reconstruction of cell-type-specific gene expression. To assess cell type composition accuracy across spots, we again use Pearson correlation between ground truth and deconvolved proportions for each matched cell type. Additionally, we calculate the root mean squared error (RMSE) for each spot to quantify the discrepancy in estimated cell type proportions. Specifically, for *K* cell types, with *θ*_*k*_ denoting the true proportion and 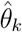 the deconvolved proportion for cell type *k*, the RMSE is computed as:

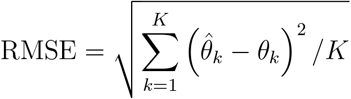

To statistically compare RMSE distributions across methods, we employ the one-sided Diebold-Mariano test to determine whether gnSPADE yields significantly lower prediction errors.

For real datasets where ground truth is unavailable, we assess performance based on how well the spatial distributions of deconvolved cell types align with histological features. For example, in the PDAC dataset, we qualitatively and spatially validate cell-type maps by examining their concordance with annotated tissue domains, enabling intuitive interpretation of spatial organization.

## 3 Simulation

### 3.1 Model-based ST data simulation

We first evaluated the performance of gnSPADE in recovering cell type mixtures and their transcriptional profiles using synthetically generated data from a model-based simulation. Unlike data-driven simulations that rely on real gene expression profiles, this synthetic setting was constructed by sampling gene frequencies across cell types and cell type proportions across spots from Dirichlet distributions[44]. This design enabled a controlled environment with known ground truth, allowing precise benchmarking of deconvolution accuracy. Specifically, we simulated *K* = 4 distinct cell types and generated pseudo-expression profiles for 100 genes conditioned on each cell type. We considered two scenarios: (1) gene co-expression patterns that are specific to a single cell type, where related genes are strongly correlated only within that type (Fig.2a) labeled as “Cell Type Speicif Related” in Fig.2, and (2) gene co-expression patterns shared across all cell types (Fig.S1a) labeled as “General Related” in Fig.2d. Cell type proportions were randomly assigned across 1,000 simulated spots (Fig.2a). The resulting count matrix was generated from a multinomial distribution, balancing computational efficiency and interpretability. While simplified, this synthetic framework is well-suited for isolating model-specific behaviors and evaluating methodological strengths. Future work will extend to data-driven simulations using full gene sets to better approximate biological complexity.

**Figure 2:**
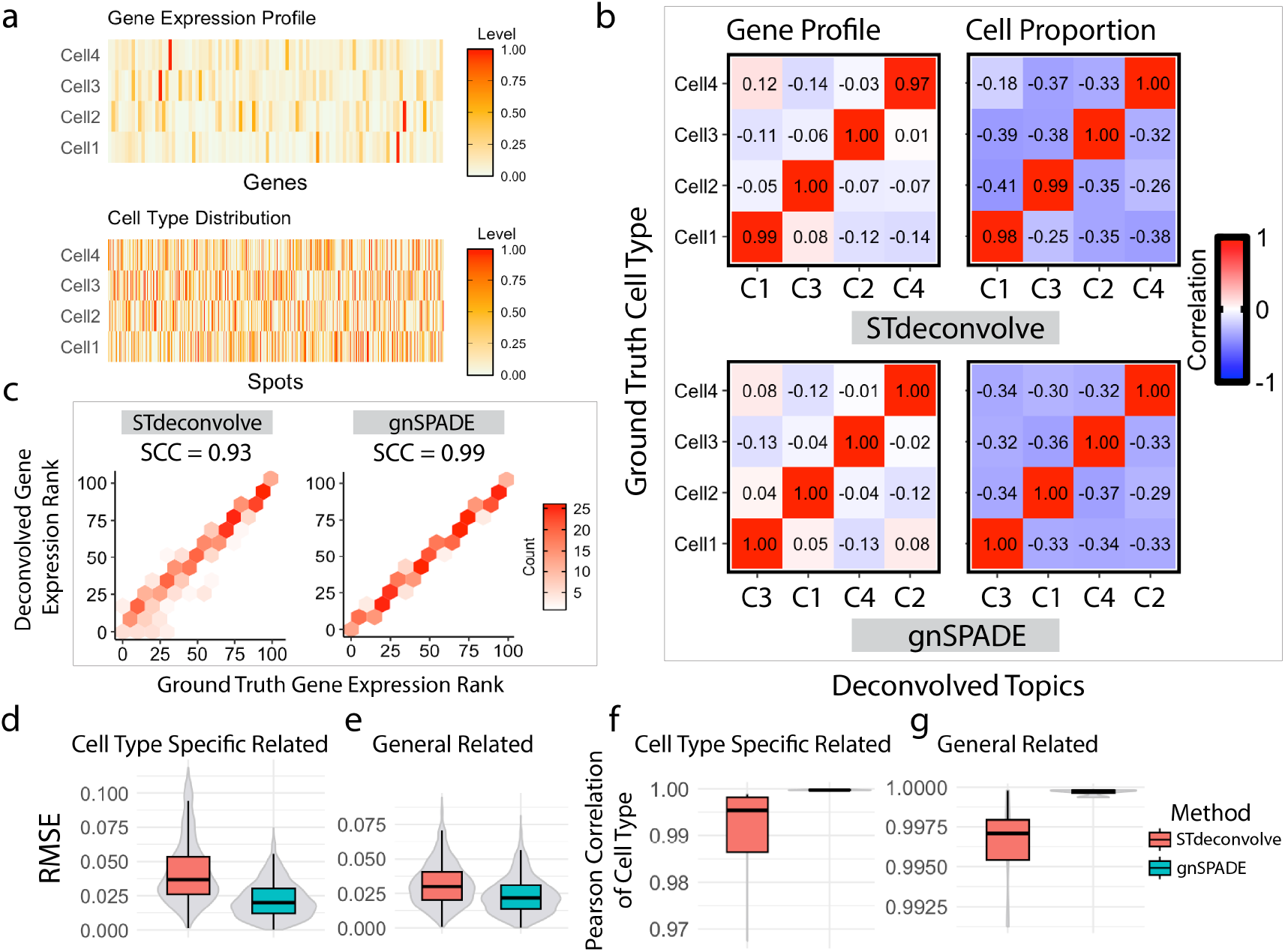
Deconvolution of Model-based ST data. **a**. Ground truth gene expression profiles for each cell type (top) and ground truth cell type proportions across all spatial spots (bottom) in the cell type-specific gene correlation simulation. **b**. PCC comparisons between ground truth and deconvolved results for the cell type-specific simulation. Top: STdeconvolve; Bottom: gnSPADE. Left: Transcriptional profiles (ground truth vs. deconvolved); Right: Cell type compositions (ground truth vs. deconvolved). **c**. Gene ranking consistency between ground truth and deconvolved cell-type transcriptional profiles. Each point represents the rank of a gene’s expression in the deconvolved profile versus its rank in the corresponding ground truth profile. The associated SCC quantifies ranking agreement. Left: STdeconvolve; Right: gnSPADE. **d, e**. Boxplots of root mean squared error (RMSE) between deconvolved and ground truth cell type proportions across all spots with **d**: cell type-specific simulation and **e**: general gene correlation simulation. **f, g**. Boxplots of PCC between deconvolved and ground truth gene expression profiles with **f**: cell type-specific simulation and **g**: general gene correlation simulation.

We applied gnSPADE and STdeconvolve to these two simulated datasets to compare their deconvolution performance. For benchmarking, we matched each deconvolved cell type to its corresponding ground truth cell type by computing Pearson correlation coefficients (PCCs) between their inferred and true gene expression profiles (Fig.2b and Fig.S1b). We also evaluated the agreement between estimated and true cell type proportions across all spots. gnSPADE consistently outperformed STdeconvolve, yielding higher correlations in both transcriptional profiles and cell type compositions. In particular, Spearman correlation coefficients (SCCs) between inferred and ground truth transcriptional profiles demonstrated gnSPADE’s superior performance (Fig.2c, Fig.S1c). Further comparisons, summarized in Fig.2d-g, show that gnSPADE achieved significantly lower RMSE in cell type proportion estimates and higher PCCs in gene expression recovery compared to STdeconvolve. Notably, gnSPADE attained a significantly lower RMSE (Diebold-Mariano test p-value *<* 2.2 *×* 10^−16^), confirming its improved precision in reconstructing both cell type-specific gene expression and spatial composition.

### 3.2 Model-free ST data simulation

To evaluate the robustness and effectiveness of gnSPADE, we simulated two ST datasets based on single-cell resolution data obtained from multiplex error-robust fluorescence in situ hybridization (MERFISH)[8]. The gene network for each dataset was constructed using STRING[28], based on the number of genes and the complexity of their interactions; further details are provided in the Supplementary Notes. We first analyzed MERFISH data from the mouse medial preoptic area (MPOA)[45], where 135 genes were spatially mapped to distinguish major non-neuronal cell types and neuronal subtypes. The dataset provides gene counts at single-cell resolution, enabling high-fidelity spatial transcriptomic profiling. In total, nine major cell types, including both excitatory and inhibitory neurons, were annotated. We selected one MPOA sample comprising 49,138 cells from 12 tissue sections of a female mouse brain. To generate pseudo-ST data, we aggregated single-cell gene expression into non-overlapping grids of 100*µm*^2^, yielding 3,072 spatial spots across all sections (256 grids per section; Fig.3a). The proportion of cell types in each grid served as the ground truth for cell type composition, while the average gene expression profile per cell type was treated as the ground truth for transcriptional profiles.

**Figure 3:**
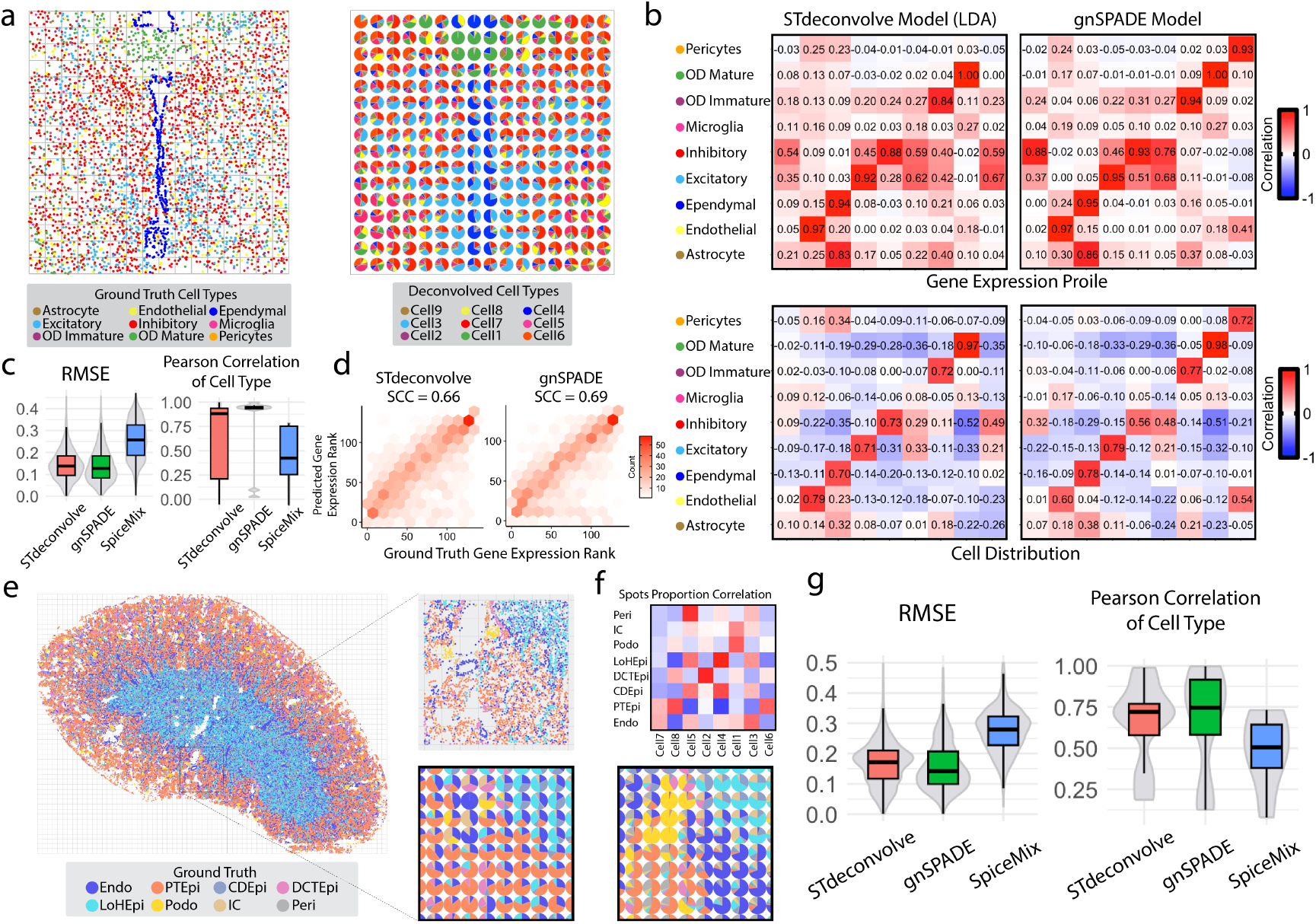
Deconvolution of MPOA (a-d) and MK (e-g) simulated ST data. **a** (Left) Ground truth single-cell resolution MERFISH data from a section of the medial preoptic area (MPOA), aggregated into 100*µm*^2^ grids (grey squares). (Right) Predicted spatial cell type composition from gnSPADE for the same region. **b** Heatmaps of Pearson correlation coefficients comparing ground truth with deconvolved results. Left: STdeconvolve; Right: gnSPADE. Top panels show correlations of transcriptional profiles; bottom panels show correlations of cell type compositions across spatial spots. **c** (Left) RMSE of deconvolved cell type proportions across spots. (Right) PCC between deconvolved transcriptional profiles and matched ground truth cell types. **d** Gene expression ranking consistency. Each gene is ranked by expression level in the deconvolved transcriptional profiles and compared to its rank in the matched ground truth profile. The corresponding SCC quantifies rank similarity. **e** (Left) Spatial layout of single-cell resolution MK MERFISH data. (Top right) Zoomed-in view of a selected region. (Bottom right) Pie chart showing aggregated cell type proportions within the selected area after merging single cells into spatial spots. **f** (Top) Heatmap of Pearson correlations between ground truth cell types and their matched deconvolved cell types from gnSPADE. (Bottom) Pie chart of gnSPADE-inferred cell type proportions for the zoomed-in region shown in **e. g** (Left) RMSE of deconvolved cell type proportions per spot. (Right) PCC between deconvolved transcriptional profiles and their matched ground truth cell types. **Abbreviations:** Endo, endothelial cell; PTEpi, epithelial cell of the proximal tubule; IC, immune cell; CDEpi, collecting duct epithelial cell; DCTEpi, distal convoluted tubule epithelial cell; LoHEpi, loop of Henle epithelial cell; Peri, pericyte; Podo, podocyte.

For this simulated data, we also compared with SpiceMix[21]. We fixed the number of deconvolved cell types at *K* = 9 for gnSPADE, STdeconvolve, and SpiceMix to match the true number of annotated cell types. Deconvolved cell types were aligned with ground truth cell types based on the highest PCC between their transcriptional profiles (Fig.3b and Fig.S2a). The spatial distribution of gnSPADE-inferred cell type proportions closely mirrored the patterns observed in the ground truth MERFISH data (Fig.3a). Compared with STdeconvolve and SpiceMix, gnSPADE achieved higher PCC values for both transcriptional profiles and cell type compositions across spots (Fig.3b), along with stronger SCC in gene expression ranking (Fig.3d). gnSPADE also produced the lowest RMSE in cell type proportion estimates relative to the ground truth across simulated spots (Fig.3c), significantly outperforming other methods (Diebold-Mariano test p-value *<* 2.2 *×* 10^−16^).

We further evaluated gnSPADE using another publicly available MERFISH single-cell dataset from the mouse kidney (MK)[7], which profiles 307 genes across 126,547 cells annotated into eight distinct cell types. Following a similar strategy as with the MPOA data, we simulated spatial transcriptomics (ST) data by partitioning the MK dataset into 2,472 spatially contiguous grids and aggregating gene expression from cells within each grid to create spot-level ST data (Fig.3e). This approach provided ground truth for both cell type compositions and transcriptional profiles, based on the average expression per cell type.

We applied gnSPADE, alongside STdeconvolve and SpiceMix, to the simulated MK ST dataset with the number of cell types fixed at eight. Deconvolved cell types were matched to the ground truth using PCC of their transcriptional profiles. gnSPADE exhibited consistently stronger correlations between deconvolved and ground truth gene expression profiles, as well as between inferred and true cell type proportions across all spots (Fig.S3a). While PCC analyses quantified the fidelity of transcriptional and compositional recovery (Fig.3f, Fig.S3a), pie chart visualizations further demonstrated that gnSPADE more accurately reproduced the spatial distribution patterns of cell types compared to STdeconvolve and SpiceMix (Fig.3f, Fig.S3b). Additionally, gnSPADE achieved the lowest RMSE in estimating cell type proportions per spot, significantly outperforming the other methods (Diebold-Mariano test p-value *<* 2.2 *×* 10^−16^). These results confirm that gnSPADE consistently delivers superior deconvolution performance, reflected by higher PCC values for transcriptional profiles(Fig.3g).

## 4 Benchmark with real ST data deconvolution

### 4.1 Mouse olfactory bulb ST deconvolution

We first focused on the mouse main olfactory bulb (MOB) ST dataset[1], which features five primary symmetric layers identified through histological analysis: the granule cell layer (GCL), the mitral cell layer (MCL), the outer plexiform layer (OPL), the glomerular layer (GL), and the olfactory nerve layer (ONL)[46] (Fig.S4a). Using gnSPADE, we identified the optimal number of deconvolved cell types to be *K* = 9 (Fig.S4b).

To evaluate deconvolution accuracy, we compared gnSPADE and STdeconvolve by correlating the inferred cell type proportions with the known spatial layer annotations of the MOB. This allowed us to align deconvolved cell types with anatomical layers and assess whether the inferred spatial distributions matched expected tissue structure. gnSPADE-predicted cell type compositions closely mirrored the layered organization of the MOB (Fig.4a). As shown in the correlation heatmap (Fig.4b), gnSPADE significantly improved alignment between deconvolved cell types and known layers, particularly in the glomerular layer (GL). Specifically, gnSPADE-identified cell type C7, enriched in the GL, exhibited a strong correlation of 0.55 with the GL region, indicating robust spatial correspondence. In contrast, the matched STdeconvolve-inferred cell type X1 showed only weak correlation with GL (0.34) and was scattered across unrelated spots, lacking clear structural alignment (Fig.4b,c).

**Figure 4:**
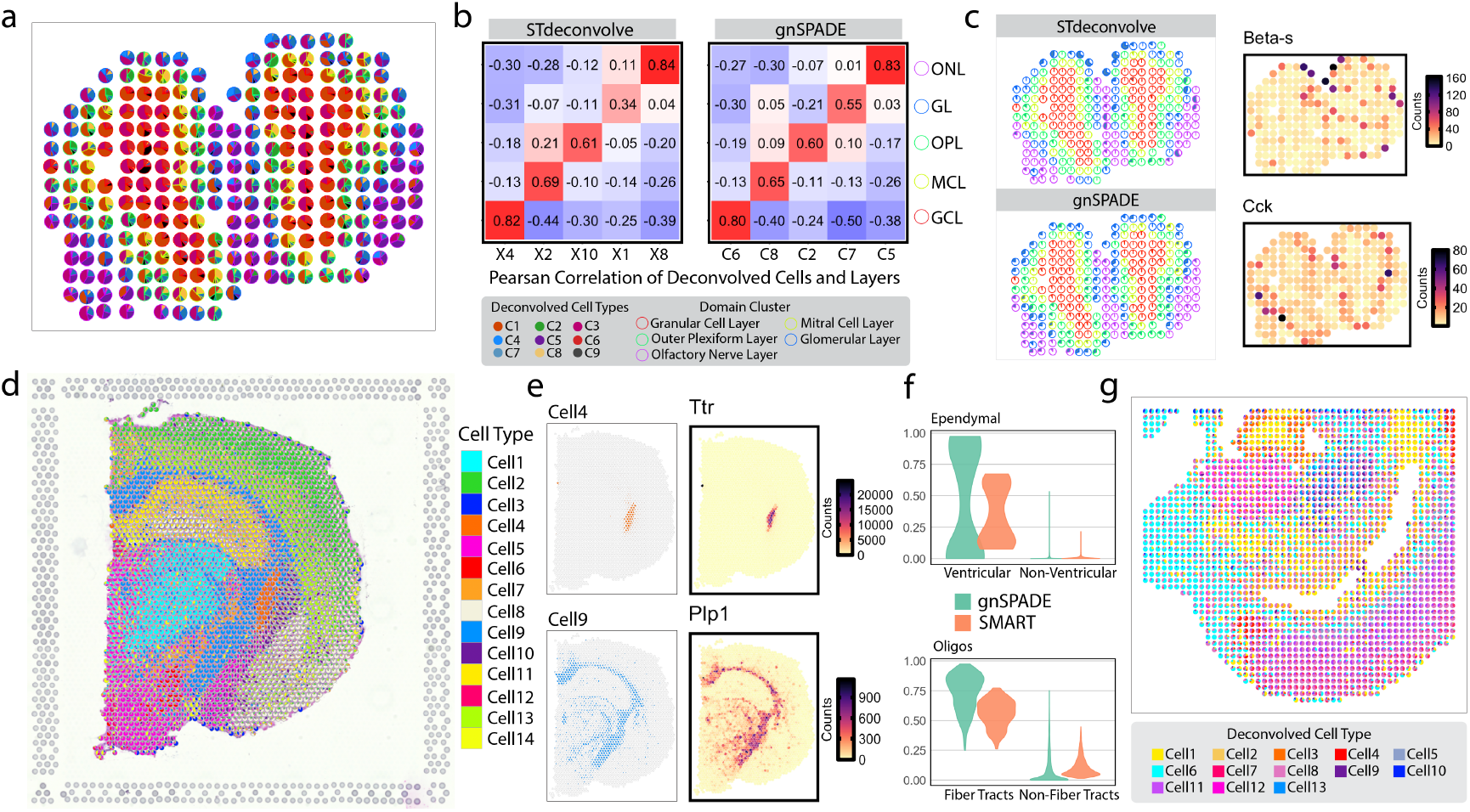
Deconvolution of mouse MOB (a-c), brain (d-f) and embryo (g) ST data. **a** gnSPADE-inferred cell type proportions for the MOB dataset, visualized as pie charts at each spot. Spot outlines are color-coded according to cluster assignments that correspond to annotated MOB layers. **b** Heatmap of PCC between known layer annotations and gnSPADE-inferred cell type proportions across spots. Strong diagonal values indicate well-aligned deconvolved cell types with anatomical layers. **c** Identification of cell types enriched in the glomerular layer (GL) and their associated top marker genes for STdeconvolve (top) and gnSPADE (bottom). Left: Pie charts show the distribution of the matched deconvolved cell type within the GL, using the same spot outline colors as in **a**. Right: Gene count distributions for the top marker gene linked to the matched deconvolved cell type in the GL for each method. **d** gnSPADE-inferred cell type proportions for mouse brain data from the 10x Genomics Visium platform, shown as pie charts per spot. **e** Left: Spatial prediction of two highlighted deconvolved cell types. Right: Corresponding expression patterns of their top differentially expressed genes across all spots (Top: ventricular; Bottom: fiber tracts). **f** Violin plots comparing the predicted cell type proportions by gnSPADE and SMART in annotated regions. Top: Ventricular region; Bottom: Fiber tracts. **g** gnSPADE-inferred cell type proportions for DBiT-seq data from the lower body of an E11 mouse embryo.

We further assessed the biological relevance of these results by examining the top-expressed genes in each deconvolved cell type. The top expressed gene in STdeconvolve’s topic X1, *Beta-s*, showed poor spatial alignment with the GL. In contrast, gnSPADE’s topic C7 was characterized by high expression of *Cck*, a known marker of the GL[47], which matched the GL structure with high spatial fidelity. These findings support the practical interpretability of gnSPADE’s deconvolution results.

Additionally, gnSPADE-identified cell type C9 overlapped with the granule cell layer previously identified via clustering and was located in a region consistent with the rostral migratory stream (RMS)[48], further validating the spatial accuracy of the inferred cell types.

### 4.2 Mouse brain and embryo ST deconvolution

We next applied gnSPADE to an ST data from a coronal section of mouse brain generated using the 10x Genomics Visium platform[49] (Fig.4d). In 10x Visium, mRNAs are captured from tissue sections using DNA-barcoded spots with a resolution of 55*µm*^2^, typically encompassing 1-10 cells per spot. gnSPADE identified the optimal number of deconvolved cell types as *K* = 14 (Fig.S5a), revealing spatially distinct patterns consistent with known brain structures from the Allen Brain Atlas[50], such as the ventricular and fiber tract regions (Fig.S5b; Fig.4e).

Differential gene expression analysis further supported the biological relevance of these deconvolved cell types. For instance, *Ttr*, a known marker of ventricular structures, and *Plp1*, a well-established marker for oligodendrocytes essential for axonal myelination [51], were identified as top genes in the corresponding deconvolved cell types. Both genes exhibited spatial expression patterns that matched expected anatomical locations, reinforcing the accuracy of gnSPADE’s spatial deconvolution (Fig.4e).

To further assess gnSPADE’s performance, we compared its results with those from SMART, a semi-supervised, marker-assisted topic modeling method that uses a curated list of cell-type-specific marker genes. While SMART directly associates deconvolved topics with known cell types, its reliance on pre-defined markers may limit flexibility in capturing biologically relevant variability. gnSPADE predicted significantly higher proportions of oligodendrocytes in fiber tract regions compared to SMART, and lower proportions in non-fiber tract regions, consistent with known oligodendrocyte localization (Fig.4f). In the ventricular region, gnSPADE again outperformed SMART in identifying region-specific cell types (Fig.4f). Although SMART annotated these cells as ependymal cells, the top marker gene *Ttr* that is highly expressed and spatially restricted to the ventricular region, is not expressed by ependymal cells but rather by the choroid plexus epithelium, which resides specifically in the lateral ventricles[52]. Ependymal cells, while lining the ventricular walls and aiding cerebrospinal fluid (CSF) circulation, do not express *Ttr*[53], underscoring the limitations of marker-only annotation approaches and highlighting the importance of examining gene expression profiles even in semi-reference based methods.

In addition to the ventricular and fiber tract regions, gnSPADE also successfully identified other key brain regions. For example, in the hippocampal region, gnSPADE highlighted *Hpcs*, a strong hippocampal marker gene[54], which showed region-specific expression consistent with anatomical expectations (Fig.S5d).

We next applied gnSPADE to spatial transcriptomics data from the lower body of an E11 mouse embryo, generated using DBiT-seq[55], which offers high-resolution profiling at 25*µm*^2^ resolution. gnSPADE identified an optimal number of cell types as *K* = 13 (Fig.S6a), consistent with the number of transcriptionally and spatially distinct regions reported by the original study[55], including the atrium, ventricle, liver, and erythrocyte-enriched blood vessels (Fig.4g; Fig.S6b). The spatial distributions of gnSPADE-deconvolved cell types closely matched these anatomical structures and aligned well with histological annotations from the original dataset. Notably, gnSPADE recovered well-defined spatial domains corresponding to key developmental features such as the ventricle, fetal liver, and neural tube (Fig.S6c). Moreover, the top differentially expressed genes for each deconvolved cell type included known markers specific to these anatomical regions, further supporting the biological validity of gnSPADE’s deconvolution results (Fig.S6d).

### 4.3 Human PDAC ST data deconvolution

We further applied gnSPADE to human pancreatic ductal adenocarcinoma (PDAC) ST data obtained from microarray slides[56]. This dataset contains multiple histologically annotated regions, including cancer, ductal, pancreatic, and stroma, based on H&E staining (Fig.5a). We evaluated gnSPADE alongside four other deconvolution methods: two reference-free methods (STdeconvolve and SpiceMix), and two marker gene-required approaches (CARD-free[14] and SMART[57]) (Fig.5b). While CARD-free is labeled as reference-free, it still requires marker genes as selected features in the count matrix and employs a conditional autoregressive non-negative matrix factorization model. However, it can only identify a fixed number of cell types specified in the input list and does not directly annotate them. To ensure a fair comparison, we used the 1,379 marker genes provided by CARD-free for all methods and set the number of deconvolved cell types to 20 to match the reference list.

**Figure 5:**
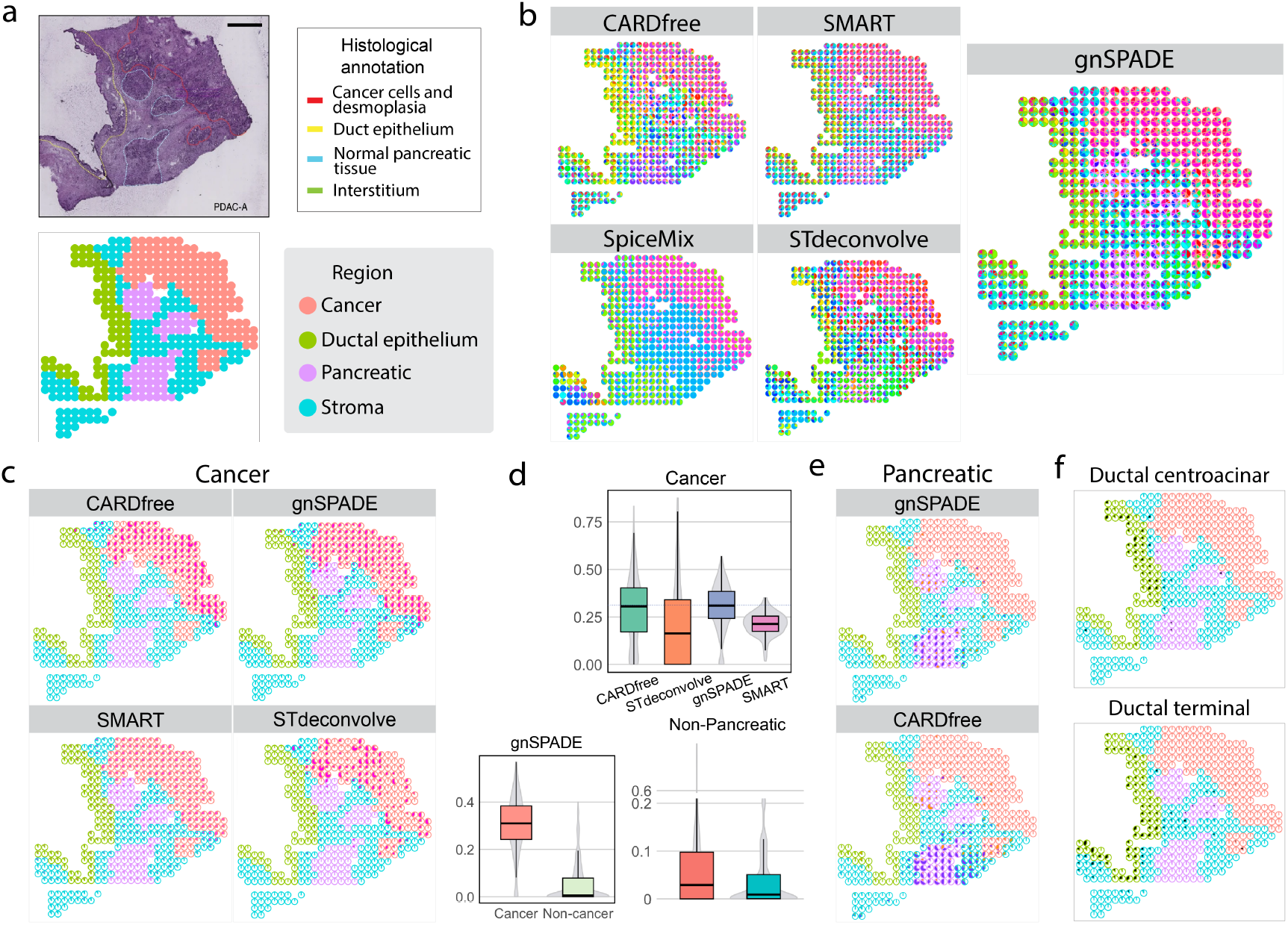
Deconvolution of the PDAC ST data. **a** (Top) Histological staining image of the tissue section. (Bottom) Spatial regions annotated based on the original study. **b** Inferred cell type compositions from five deconvolution methods, CARD-free, SMART, SpiceMix, STdeconvolve, and gnSPADE, visualized as pie charts per spot. Spots are outlined according to the annotated regions shown in **a**. gnSPADE results (far right) exhibit spatial patterns that more closely align with the ground truth annotations. **c** Highlights of deconvolved tumor-related cell types identified by CARD-free, SMART, STdeconvolve, and gnSPADE, shown as spot-level pie charts. **d** (Top) Boxplot comparing the proportions of tumor-related deconvolved cell types within cancer-annotated regions. (Bottom left) Comparison of gnSPADE-inferred tumor cell type proportions between cancerous and non-cancerous regions. (Bottom right) Comparison of gnSPADE and CARD-free in predicting cell type proportions within non-pancreatic regions. **e** Spatial distribution of predicted pancreatic region-related sub-cell types. Top: gnSPADE; Bottom: CARD-free. **f** Spatial distribution of ductal sub-cell types inferred by gnSPADE.

gnSPADE outperformed all competing methods in spatial accuracy and biological interpretability. In cancer regions, gnSPADE produced the clear spatial patterns and predicted significantly higher proportions of cancer cell populations compared to non-cancerous regions (t-test p-value = 2.2 *×* 10^−16^) (Fig.5c,d). While SpiceMix also identified high proportions of cancer cells, it incorrectly assigned cancer-like patterns to non-cancerous spots, likely due to its reliance on spatial continuity (Fig.S7a). SpiceMix also underperformed in regions lacking clear spatial structure, where it failed to distinguish biologically distinct regions (Fig.S7b).

gnSPADE further revealed nuanced cellular heterogeneity in non-cancerous regions through top gene analysis. In the pancreatic region, gnSPADE identified transcriptionally distinct acinar-adjacent subtypes (Fig.5e), which shared common markers but diverged in pathway-level activity (Fig.S7c,d). Reactome pathway analysis of Cell3 and Cell10 revealed canonical exocrine functions, such as protein digestion and lipid metabolism, in both, consistent with acinar identity[58]. However, Cell10 also showed strong enrichment in immune and barrier-defense pathways, including *REG3A*/*REG3G*-driven responses, alpha-defensins, and antimicrobial peptides. This suggests that Cell3 represents a classical digestive acinar phenotype, whereas Cell10 reflects a stress-adapted subpopulation involved in innate immunity[59]. Notably, while CARD-free also detected pancreatic-related cells (Fig.5e), their spatial distribution overlapped excessively with neighboring stromal regions. gnSPADE, in contrast, showed significantly lower predicted proportions of pancreatic-related cells in non-pancreatic regions (t-test p-value = 2.2 *×* 10^−16^) (Fig.5d).

In the ductal region, gnSPADE successfully identified two distinct ductal subtypes, centroacinar and terminal ductal cells, based on the top differentially expressed genes (Fig.5f; Fig.S8a,b). Cell4 (ductal centroacinar) was enriched for pathways related to oxidative stress, wounding, and AP-1 transcriptional activity, with marker genes such as *JUN, HIF1A*, and *LITAF* suggesting active responses to hypoxia, inflammation, and epithelial remodeling (Fig.S8c). These features align with progenitor-like or transitional states observed in acinar-to-ductal metaplasia (ADM) or early PDAC[60, 61]. In contrast, Cell6 (ductal terminal) expressed hallmark genes of secretory ductal cells, *TFF* family genes, *MUC5B, PIGR, SLC7A5, CES1*, and *CRISP3*, implicated in mucosal protection, immune crosstalk, and epithelial barrier maintenance. Pathway enrichment revealed strong representation of functions related to stress response, transcellular transport, and epithelial organization, consistent with a terminal ductal phenotype. These transcriptional profiles resemble goblet-like or salivary gland-like ductal epithelia described in human pancreas and salivary tissues, particularly under inflammatory or metaplastic conditions[62, 63].

## 5 Discussion

Understanding the cellular composition within each spatial location is essential for interpreting ST data at multicellular resolution. gnSPADE addresses this need with a reference-free, probabilistic framework that infers cell-type-specific gene expression and cell type composition directly from the data, without relying on predefined marker genes or external single-cell references. This makes it particularly valuable for studying novel tissues or poorly characterized disease contexts where reference data are incomplete or unavailable. gnSPADE can also be used in follow-up analyses to further resolve substructures within purified cell populations, enabling the discovery of subtle heterogeneity and latent sub-cell types.

Across both simulated and real datasets, gnSPADE consistently demonstrated superior performance compared to existing reference-free methods. In simulations, whether model-generated data or derived from high-resolution single-cell data like MERFISH, gnSPADE outperformed STdeconvolve and SpiceMix, identifying spatially coherent cell types with lower reconstruction error and stronger correlation to ground truth. In real ST applications, gnSPADE achieved better alignment with histologically defined regions across diverse platforms and resolutions, including the mouse olfactory bulb, embryo, coronal section of the mouse brain, and PDAC samples. Compared to semi-supervised methods such as SMART and CARD-free, gnSPADE showed greater robustness and interpretability. SMART depends on accurate marker gene lists to identify known cell types, while CARD-free requires a fixed number of input cell types and relies on marker genes as feature filters. Both are sensitive to incomplete or noisy marker gene sets, which could lead to ambiguous or biologically inconsistent deconvolution outcomes.

In the PDAC analysis, gnSPADE provided more spatially accurate predictions of cancer, pancreatic, and ductal subtypes compared to other methods. While SpiceMix was able to recover high cancer cell proportions due to spatial continuity modeling, it misclassified regions lacking such continuity, highlighting its limitations in irregular tissue contexts. Moreover, when overestimating the number of cell types, SpiceMix’s inferred transcriptional profiles became overly sparse, reducing interpretability[21].

In contrast, gnSPADE operates as a fully unsupervised generative model. Its integration of gene-gene correlation information addresses a key limitation of STdeconvolve, which assumes gene independence. By leveraging network-informed priors via a Markov random field, gnSPADE improves the coherence and biological interpretability of deconvolved cell types. This enables not only more accurate estimation of cell type proportions and transcriptional profiles across platforms, but also enhances our understanding of spatial tissue architecture, cellular neighborhoods, and potential cell-cell interactions relevant to development and disease.

Despite these advantages, gnSPADE, like other LDA-based approaches, has limitations. Its performance is influenced by dataset characteristics, such as the number of spots and genes, and is sensitive to gene selection strategies[64]. Additionally, gnSPADE does not currently incorporate spatial priors and treats spots as independent observations. Future work could enhance model accuracy by integrating spatial relationships and scaling inference through more computationally efficient algorithms.

In summary, gnSPADE represents a powerful and flexible reference-free deconvolution framework for ST data. By incorporating gene network structures, it outperforms existing reference-free approaches and performs comparably to semi-supervised methods, without requiring external annotations. gnSPADE is well-suited for unsupervised clustering, cell subtype identification, and spatial data visualization, offering a valuable tool for dissecting tissue complexity and gene expression variation across diverse biological contexts.

## Author contribution

Y.C. conceived the idea. A.X. and Y.C. designed the experiments. A.X. developed the method, implemented the software, performed simulations, and analyzed real data. A.X. and Y.C. wrote the manuscript.

## Conflict of interest

None declared.

## Data availability

The MERFISH dataset of the mouse medial preoptic area[45] is available for download from https://datadryad.org/stash/dataset/doi:10.5061/dryad.8t8s248/. The MK dataset[7] are available for download from https://figshare.com/projects/MERFISH_mouse_comparison_study/134213. Data and H&E images for all MOB replicates[1] are available for download at https://www.spatialresearch.org/resources-published-datasets/doi-10-1126science-aaf2403/. The 10x Visium coronal section of the mouse brain dataset[49] is available for download at https://www.10xgenomics.com/resources/datasets/mouse-brain-section-coronal-1-standard-1-1-0. DBiT-seq dataset of E11 mouse embryo lower body (GSM4364242 E11-1L)[55] is available for download at https://www.ncbi.nlm.nih.gov/geo/query/acc.cgi?acc=GSE137986. The PDAC data were obtained from sample A of the PDAC dataset and are available at the Gene Expression Omnibus (accession number GSE111672)[56].

## Code availability

R codes to implement gnSPADE are available in the GitHub repository at https://github.com/Cui-STT-Lab/gnSPADE

## Notes

### Competing Interest Statement

The authors have declared no competing interest.

## References

[1] P. L. Ståhl, F. Salmén, S. Vickovic, A. Lundmark, J. F. Navarro, J. Magnusson, S. Giacomello, M. Asp, J. O. Westholm, M. Huss, et al., “Visualization and analysis of gene expression in tissue sections by spatial transcriptomics,” Science, vol. 353, no. 6294, pp. 78–82, 2016.

[2] D. J. Burgess, “Spatial transcriptomics coming of age,” Nature Reviews Genetics, vol. 20, no. 6, pp. 317–317, 2019.

[3] L. Larsson, J. Frisén, and J. Lundeberg, “Spatially resolved transcriptomics adds a new dimension to genomics,” Nature Methods, vol. 18, no. 1, pp. 15–18, 2021.

[4] V. Svensson, S. A. Teichmann, and O. Stegle, “Spatialde: identification of spatially variable genes,” Nature Methods, vol. 15, no. 5, pp. 343–346, 2018.

[5] S. Sun, J. Zhu, and X. Zhou, “Statistical analysis of spatial expression patterns for spatially resolved transcriptomic studies,” Nature Methods, vol. 17, no. 2, pp. 193–200, 2020.

[6] A. Janesick, R. Shelansky, A. D. Gottscho, F. Wagner, S. R. Williams, M. Rouault, G. Beliakoff, C. A. Morrison, M. F. Oliveira, J. T. Sicherman, et al., “High resolution mapping of the tumor microenvironment using integrated single-cell, spatial and in situ analysis,” Nature Communications, vol. 14, no. 1, p. 8353, 2023.

[7] J. Liu, V. Tran, V. N. P. Vemuri, A. Byrne, M. Borja, Y. J. Kim, S. Agarwal, R. Wang, K. Awayan, A. Murti, et al., “Concordance of merfish spatial transcriptomics with bulk and single-cell rna sequencing,” Life Science Alliance, vol. 6, no. 1, 2023.

[8] K. H. Chen, A. N. Boettiger, J. R. Moffitt, S. Wang, and X. Zhuang, “Spatially resolved, highly multiplexed rna profiling in single cells,” Science, vol. 348, no. 6233, p. aaa6090, 2015.

[9] V. Marx, “Method of the year: spatially resolved transcriptomics,” Nature Methods, vol. 18, no. 1, pp. 9–14, 2021.

[10] H. Su, Y. Wu, B. Chen, and Y. Cui, “Stance: a unified statistical model to detect cell-type-specific spatially variable genes in spatial transcriptomics,” Nature Communications, vol. 16, no. 1, p. 1793, 2025.

[11] D. M. Cable, E. Murray, L. S. Zou, A. Goeva, E. Z. Macosko, F. Chen, and R. A. Irizarry, “Robust decomposition of cell type mixtures in spatial transcriptomics,” Nature Biotechnology, vol. 40, no. 4, pp. 517–526, 2022.

[12] V. Kleshchevnikov, A. Shmatko, E. Dann, A. Aivazidis, H. W. King, T. Li, R. Elmentaite, A. Lomakin, V. Kedlian, A. Gayoso, et al., “Cell2location maps fine-grained cell types in spatial transcriptomics,” Nature Biotechnology, vol. 40, no. 5, pp. 661–671, 2022.

[13] M. Elosua-Bayes, P. Nieto, E. Mereu, I. Gut, and H. Heyn, “Spotlight: seeded nmf regression to deconvolute spatial transcriptomics spots with single-cell transcriptomes,” Nucleic Acids Research, vol. 49, no. 9, pp. e50–e50, 2021.

[14] Y. Ma and X. Zhou, “Spatially informed cell-type deconvolution for spatial transcriptomics,” Nature Biotechnology, vol. 40, no. 9, pp. 1349–1359, 2022.

[15] H. Li, J. Zhou, Z. Li, S. Chen, X. Liao, B. Zhang, R. Zhang, Y. Wang, S. Sun, and X. Gao, “A comprehensive benchmarking with practical guidelines for cellular deconvolution of spatial transcriptomics,” Nature Communications, vol. 14, no. 1, p. 1548, 2023.

[16] L. Yan and X. Sun, “Benchmarking and integration of methods for deconvoluting spatial transcriptomic data,” Bioinformatics, vol. 39, no. 1, p. btac805, 2023.

[17] A. Kiemen, A. M. Braxton, M. P. Grahn, K. S. Han, J. M. Babu, R. Reichel, F. Amoa, S.-M. Hong, T. C. Cornish, E. D. Thompson, et al., “In situ characterization of the 3d microanatomy of the pancreas and pancreatic cancer at single cell resolution,” BioRxiv, pp. 2020–12, 2020.

[18] Q. H. Nguyen, N. Pervolarakis, K. Nee, and K. Kessenbrock, “Experimental considerations for single-cell rna sequencing approaches,” Frontiers in Cell and Developmental Biology, vol. 6, p. 108, 2018.

[19] A. Haque, J. Engel, S. A. Teichmann, and T. Lönnberg, “A practical guide to single-cell rna-sequencing for biomedical research and clinical applications,” Genome Medicine, vol. 9, pp. 1–12, 2017.

[20] A. Deutsch, D. Feng, J. E. Pessin, and K. Shinoda, “The impact of single-cell genomics on adipose tissue research,” International Journal of Molecular Sciences, vol. 21, no. 13, p. 4773, 2020.

[21] B. Chidester, T. Zhou, S. Alam, and J. Ma, “Spicemix enables integrative single-cell spatial modeling of cell identity,” Nature Genetics, vol. 55, no. 1, pp. 78–88, 2023.

[22] B. F. Miller, F. Huang, L. Atta, A. Sahoo, and J. Fan, “Reference-free cell type deconvolution of multi-cellular pixel-resolution spatially resolved transcriptomics data,” Nature Communications, vol. 13, no. 1, p. 2339, 2022.

[23] D. M. Blei, A. Y. Ng, and M. I. Jordan, “Latent dirichlet allocation,” Journal of Machine Learning Research, vol. 3, no. Jan, pp. 993–1022, 2003.

[24] T. Stuart and R. Satija, “Integrative single-cell analysis,” Nature Reviews Genetics, vol. 20, no. 5, pp. 257–272, 2019.

[25] B. Zhang and S. Horvath, “A general framework for weighted gene co-expression network analysis,” Statistical Applications in Genetics and Molecular Biology, vol. 4, no. 1, 2005.

[26] S. Aibar, C.B. González-Blas, T. Moerman, V. A. Huynh-Thu, H. Imrichova, G. Hulselmans, F. Rambow, J.-C. Marine, P. Geurts, J. Aerts, et al., “Scenic: single-cell regulatory network inference and clustering,” Nature Methods, vol. 14, no. 11, pp. 1083–1086, 2017.

[27] P. Xie, D. Yang, and E. Xing, “Incorporating word correlation knowledge into topic modeling,” in Proceedings of the 2015 conference of the north American chapter of the association for computational linguistics: human language technologies, pp. 725–734, 2015.

[28] D. Szklarczyk, A. L. Gable, K. C. Nastou, D. Lyon, R. Kirsch, S. Pyysalo, N. T. Doncheva, M. Legeay, T. Fang, P. Bork, et al., “The string database in 2021: customizable protein– protein networks, and functional characterization of user-uploaded gene/measurement sets,” Nucleic Acids Research, vol. 49, no. D1, pp. D605–D612, 2021.

[29] V. A. Huynh-Thu, A. Irrthum, L. Wehenkel, and P. Geurts, “Inferring regulatory networks from expression data using tree-based methods,” PloS One, vol. 5, no. 9, p. e12776, 2010.

[30] M. Kanehisa, M. Furumichi, Y. Sato, M. Ishiguro-Watanabe, and M. Tanabe, “Kegg: integrating viruses and cellular organisms,” Nucleic Acids Research, vol. 49, no. D1, pp. D545–D551, 2021.

[31] A. Fabregat, S. Jupe, L. Matthews, K. Sidiropoulos, M. Gillespie, P. Garapati, R. Haw, B. Jassal, F. Korninger, B. May, et al., “The reactome pathway knowledgebase,” Nucleic Acids Research, vol. 46, no. D1, pp. D649–D655, 2018.

[32] T. L. Griffiths and M. Steyvers, “Finding scientific topics,” Proceedings of the National Academy of Sciences, vol. 101, no. suppl 1, pp. 5228–5235, 2004.

[33] D. M. Blei, “Mixed-membership models (and an introduction to variational inference),” Course notes for Foundations of Graphical Models. Nov, 2015.

[34] N. Baumann and D. Pham-Dinh, “Biology of oligodendrocyte and myelin in the mammalian central nervous system,” Physiological Reviews, vol. 81, no. 2, pp. 871–927, 2001.

[35] O. Jahn, S. Tenzer, and H. B. Werner, “Myelin proteomics: molecular anatomy of an insulating sheath,” Molecular Neurobiology, vol. 40, no. 1, pp. 55–72, 2009.

[36] J. Qiang, P. Chen, T. Wang, and X. Wu, “Topic modeling over short texts by incorporating word embeddings,” in Advances in Knowledge Discovery and Data Mining: 21st Pacific-Asia Conference, PAKDD 2017, Jeju, South Korea, May 23-26, 2017, Proceedings, Part II 21, pp. 363–374, Springer, 2017.

[37] X. Zhang, Y. Lan, J. Xu, F. Quan, E. Zhao, C. Deng, T. Luo, L. Xu, G. Liao, M. Yan, et al., “Cellmarker: a manually curated resource of cell markers in human and mouse,” Nucleic Acids Research, vol. 47, no. D1, pp. D721–D728, 2019.

[38] J. Fan, N. Salathia, R. Liu, G. E. Kaeser, Y. C. Yung, J. L. Herman, F. Kaper, J.-B. Fan, K. Zhang, J. Chun, et al., “Characterizing transcriptional heterogeneity through pathway and gene set overdispersion analysis,” Nature Methods, vol. 13, no. 3, pp. 241–244, 2016.

[39] L. Cowen, T. Ideker, B. J. Raphael, and R. Sharan, “Network propagation: a universal amplifier of genetic associations,” Nature Reviews Genetics, vol. 18, no. 9, pp. 551–562, 2017.

[40] R. Elyanow, B. Dumitrascu, B. E. Engelhardt, and B. J. Raphael, “netnmf-sc: leveraging gene–gene interactions for imputation and dimensionality reduction in single-cell expression analysis,” Genome Research, vol. 30, no. 2, pp. 195–204, 2020.

[41] F. Yuan, Y.-H. Zhang, S. Wan, S. Wang, and X.-Y. Kong, “Mining for candidate genes related to pancreatic cancer using protein-protein interactions and a shortest path approach,” BioMed Research International, vol. 2015, no. 1, p. 623121, 2015.

[42] E. I. Boyle, S. Weng, J. Gollub, H. Jin, D. Botstein, J. M. Cherry, and G. Sherlock, “Go:: Termfinder—open source software for accessing gene ontology information and finding significantly enriched gene ontology terms associated with a list of genes,” Bioinformatics, vol. 20, no. 18, pp. 3710–3715, 2004.

[43] T. Wu, E. Hu, S. Xu, M. Chen, P. Guo, Z. Dai, T. Feng, L. Zhou, W. Tang, L. Zhan, et al., “clusterprofiler 4.0: A universal enrichment tool for interpreting omics data,” The Innovation, vol. 2, no. 3, 2021.

[44] X. Wu, H. Wu, and Z. Wu, “Penalized latent dirichlet allocation model in single-cell rna sequencing,” Statistics in Biosciences, pp. 1–20, 2021.

[45] J. R. Moffitt, D. Bambah-Mukku, S. W. Eichhorn, E. Vaughn, K. Shekhar, J. D. Perez, N. D. Rubinstein, J. Hao, A. Regev, C. Dulac, et al., “Molecular, spatial, and functional single-cell profiling of the hypothalamic preoptic region,” Science, vol. 362, no. 6416, p. eaau5324, 2018.

[46] S. Nagayama, R. Homma, and F. Imamura, “Neuronal organization of olfactory bulb circuits,” Frontiers in Neural Circuits, vol. 8, p. 98, 2014.

[47] K. B. Seroogy, N. Brecha, and C. Gall, “Distribution of cholecystokinin-like immunoreactivity in the rat main olfactory bulb,” Journal of Comparative Neurology, vol. 239, no. 4, pp. 373–383, 1985.

[48] H. Hintiryan, L. Gou, B. Zingg, S. Yamashita, H. M. Lyden, M. Y. Song, A. K. Grewal, X. Zhang, A. W. Toga, and H.-W. Dong, “Comprehensive connectivity of the mouse main olfactory bulb: analysis and online digital atlas,” Frontiers in Neuroanatomy, vol. 6, p. 30, 2012.

[49] N. Rao, S. Clark, and O. Habern, “Bridging genomics and tissue pathology: 10x genomics explores new frontiers with the visium spatial gene expression solution,” Genetic Engineering & Biotechnology News, vol. 40, no. 2, pp. 50–51, 2020.

[50] E. S. Lein, M. J. Hawrylycz, N. Ao, M. Ayres, A. Bensinger, A. Bernard, A. F. Boe, M. S. Boguski, K. S. Brockway, E. J. Byrnes, et al., “Genome-wide atlas of gene expression in the adult mouse brain,” Nature, vol. 445, no. 7124, pp. 168–176, 2007.

[51] K. A. Lüders, J. Patzig, M. Simons, K.-A. Nave, and H. B. Werner, “Genetic dissection of oligodendroglial and neuronal plp1 function in a novel mouse model of spastic paraplegia type 2,” Glia, vol. 65, no. 11, pp. 1762–1776, 2017.

[52] S. A. Liddelow, “Development of the choroid plexus and blood-csf barrier,” Frontiers in Neuroscience, vol. 9, p. 32, 2015.

[53] A. Zeisel, H. Hochgerner, P. Lönnerberg, A. Johnsson, F. Memic, J. Van Der Zwan, M. Häring, E. Braun, L. E. Borm, G. La Manno, et al., “Molecular architecture of the mouse nervous system,” Cell, vol. 174, no. 4, pp. 999–1014, 2018.

[54] R. Coras, F. A. Siebzehnrubl, E. Pauli, H. B. Huttner, M. Njunting, K. Kobow, C. Villmann, E. Hahnen, W. Neuhuber, D. Weigel, et al., “Low proliferation and differentiation capacities of adult hippocampal stem cells correlate with memory dysfunction in humans,” Brain, vol. 133, no. 11, pp. 3359–3372, 2010.

[55] Y. Liu, M. Yang, Y. Deng, G. Su, A. Enninful, C. C. Guo, T. Tebaldi, D. Zhang, D. Kim, Z. Bai, et al., “High-spatial-resolution multi-omics sequencing via deterministic barcoding in tissue,” Cell, vol. 183, no. 6, pp. 1665–1681, 2020.

[56] R. Moncada, D. Barkley, F. Wagner, M. Chiodin, J. C. Devlin, M. Baron, C. H. Hajdu, D. M. Simeone, and I. Yanai, “Integrating microarray-based spatial transcriptomics and single-cell rna-seq reveals tissue architecture in pancreatic ductal adenocarcinomas,” Nature Biotechnology, vol. 38, no. 3, pp. 333–342, 2020.

[57] C. X. Yang, D. D. Sin, and R. T. Ng, “Smart: spatial transcriptomics deconvolution using marker-gene-assisted topic model,” Genome Biology, vol. 25, no. 1, pp. 1–21, 2024.

[58] Y. J. Wang, J. Schug, K.-J. Won, C. Liu, A. Naji, D. Avrahami, M. L. Golson, and K. H. Kaestner, “Single-cell transcriptomics of the human endocrine pancreas,” Diabetes, vol. 65, no. 10, pp. 3028–3038, 2016.

[59] G.-Y. Liou, H. Döppler, B. Necela, M. Krishna, H. C. Crawford, M. Raimondo, and P. Storz, “Macrophage-secreted cytokines drive pancreatic acinar-to-ductal metaplasia through nf-κb and mmps,” Journal of Cell Biology, vol. 202, no. 3, pp. 563–577, 2013.

[60] H. Lin, J.-F. Huang, J.-R. Qiu, H.-L. Zhang, X.-J. Tang, H. Li, C.-J. Wang, Z.-C. Wang, Z.-Q. Feng, and J. Zhu, “Significantly upregulated tacstd2 and cyclin d1 correlate with poor prognosis of invasive ductal breast cancer,” Experimental and Molecular Pathology, vol. 94, no. 1, pp. 73–78, 2013.

[61] Y. Xia, B. Li, N. Gao, H. Xia, Y. Men, Y. Liu, Z. Liu, Q. Chen, and L. Li, “Expression of tumor-associated calcium signal transducer 2 in patients with salivary adenoid cystic carcinoma: Correlation with clinicopathological features and prognosis,” Oncology Letters, vol. 8, no. 4, pp. 1670–1674, 2014.

[62] M. M. F. Qadir, S. Álvarez-Cubela, D. Klein, J. van Dijk, R. Muníz-Anquela, Y.B. Moreno-Hernández, G. Lanzoni, S. Sadiq, B. Navarro-Rubio, M.T. García, et al., “Single-cell resolution analysis of the human pancreatic ductal progenitor cell niche,” Proceedings of the National Academy of Sciences, vol. 117, no. 20, pp. 10876–10887, 2020.

[63] J. Peng, B.-F. Sun, C.-Y. Chen, J.-Y. Zhou, Y.-S. Chen, H. Chen, L. Liu, D. Huang, J. Jiang, G.-S. Cui, et al., “Single-cell rna-seq highlights intra-tumoral heterogeneity and malignant progression in pancreatic ductal adenocarcinoma,” Cell Research, vol. 29, no. 9, pp. 725–738, 2019.

[64] J. Tang, Z. Meng, X. Nguyen, Q. Mei, and M. Zhang, “Understanding the limiting factors of topic modeling via posterior contraction analysis,” in International Conference on Machine Learning, pp. 190–198, PMLR, 2014.

