## Supplementary File for "gnSPADE: a reference-free deconvolution method incorporating gene network structures in spatial transcriptomics"

### Supplementary material for "gnSPADE: a reference-free deconvolution method incorporating gene network structures in spatial transcriptomics"

Aoqi Xie and Yuehua Cui*

*Department of Statistics and Probability, Michigan State University, East Lansing, MI.*

#### 1 Deconvolution with the collapsed Gibbs sampler

Latent Dirichlet Allocation (LDA) has been proposed to identify latent topics for a given set of documents[1]. In the specific context of spatial transcriptomics, this model can be analogously applied[2]. The posterior distribution of latent variables based on the observed gene expression data can be obtained as,

$$p(\boldsymbol{\theta}, \mathbf{z}|\boldsymbol{w}, \alpha, \boldsymbol{\beta}) = \frac{p(\boldsymbol{\theta}, \mathbf{z}, \boldsymbol{w}|\alpha, \boldsymbol{\beta})}{p(\boldsymbol{w}|\alpha, \boldsymbol{\beta})} = \prod_{d=1}^{D} p(\theta_d, \mathbf{z}|\boldsymbol{w}, \alpha, \boldsymbol{\beta}). \tag{1}$$

where $w$ is the collection of the corresponding genes in the gene expression matrix, $\mathbf{z}$ is the collection of assigned cell types of corresponding genes, $\theta$ is the distribution of cell types in all spots, $\theta_d$ is the distribution of cell types in spot $d$, $\alpha$ is the uniform Dirichlet scale parameter for $\boldsymbol{\theta}$, $\boldsymbol{\beta}$ is the matrix of relative gene expressions for all cell types. $\boldsymbol{\beta}$ and $\boldsymbol{\theta}$ can be estimated with two popular methods: the Variational EM algorithm and the collapsed Gibbs sampler. In our LDA model with MRF, we implemented the latter one. Next, we briefly introduce the algorithms.

We assume that the probability of each gene in cell type $k$, denoted as $\beta_k$, follows a uniform Dirichlet distribution with scaling parameter $\eta$, i.e. $\beta_k \sim \text{Dir}(\eta)$. Then, to infer the latent parameters $\boldsymbol{\theta}$ and $\mathbf{z}$, the posterior distribution conditioned on the observed gene expression data can be obtained as:

$$p(\boldsymbol{\theta}, \mathbf{z}, \boldsymbol{\beta}|\boldsymbol{w}, \alpha, \eta) = \frac{p(\boldsymbol{\theta}, \mathbf{z}, \boldsymbol{w}, \boldsymbol{\beta}|\alpha, \eta)}{p(\boldsymbol{w}|\alpha, \eta)} = \prod_{d=1}^{D}\prod_{g=1}^{V} p(\theta_d, \mathbf{z}, \beta_k|\boldsymbol{w}, \alpha, \eta) \tag{2}$$

Next, we first show the basic Gibbs sampler for LDA model by calculating the complete conditional distributions as follows:

- The conditional distribution of cell type (component) proportions $\theta_d$ is given as

$$\begin{aligned}
p(\theta_d|\mathbf{z}, \theta_{-d}, w, \beta) &= p(\theta_d|z_d, \alpha) \propto \prod_{n=1}^{N_d} p(z_{d,n}|\theta_d)p(\theta_d|\alpha) \\
&\propto \prod_{n=1}^{N_d}\prod_{k=1}^{K} \theta_{d,k}^{z_{d,n}^k} \prod_{k=1}^{K} \theta_{d,k}^{\alpha-1} \\
&= \prod_{k=1}^{K} \theta_{d,k}^{\alpha-1+\sum_{n=1}^{N_d} z_{d,n}^k} \\
&= \mathrm{Dir}(\alpha + \sum_{n=1}^{N_d} z_{d,n}) \\
&= \mathrm{Dir}(\alpha + \mathbf{z}_d)
\end{aligned}$$

- The conditional distribution of gene expression $\beta_k$ is given as

$$\begin{aligned}
p(\beta_k|\mathbf{z}, \theta, w, \beta_{-k}) &= p(\beta_k|\mathbf{z}, w, \eta) \propto \prod_{d=1}^{D}\prod_{n=1}^{N_d} p(w_{d,n}|\beta_k)^{z_{d,n}} p(\beta_d|\eta) \\
&\propto \prod_{d=1}^{D}\prod_{n=1}^{N_d}\prod_{g=1}^{V} \beta_{k,g}^{w_{d,n}^g z_{d,n}^k} \prod_{g=1}^{V} \beta_{k,g}^{\eta-1} \\
&= \prod_{g=1}^{V} \beta_{k,g}^{\eta-1+\sum_{d=1}^{D}\sum_{n=1}^{N_d} w_{d,n}^g z_{d,n}^k} \\
&= \mathrm{Dir}(\eta + \sum_{d=1}^{D}\sum_{n=1}^{N_d} w_{d,n}^g z_{d,n}^k)
\end{aligned}$$

- The conditional distribution of cell type (component) assignment $z_{d,n}$ is given as

$$\begin{aligned}
p(z_{d,n} = k|z_{-(d,n)}, \theta, \beta, \boldsymbol{w}) &= p(z_{d,n}|\theta_{d,}, \beta, w_{d,n}) \\
&\propto p(\theta_d)p(z_{d,n} = k|\theta_d)p(w_{d,n}|\beta_k) \\
&= \theta_{d,k} p(w_{d,n}|\beta_k)
\end{aligned}$$

Gibbs Sampler exhibits a slow convergence rate. Moreover, in each iteration, if a gene expression $\beta_k$ assigns a probability of 0 to a gene $w_{d,n}$, then it will have a posterior probability of 0 under $z_{d,n}$. Consequently, the information of $z_{d,n}$ will be lost in the next iteration of $\theta_d$. To address these challenges, the Collapsed Gibbs Sampler was proposed[3][4]. The conditional probability of cell type assignment $k$ is proportional to the joint probability of the assignment and the gene:

$$p(z_{d,n} = k|z_{-(d,n)}, \boldsymbol{w}) \propto p(z_{d,n} = k, w_{d,n}|z_{-(d,n)}, w_{-(d,n)})$$

Given the cell type proportions and gene expressions, the joint distribution of a cell type assignment and genes is:

$$\begin{aligned}
p(z_{d,n} = k, w_{d,n}|\theta_d, \beta_{1:K}) &= p(z_{d,n} = k|\theta_d)p(w_{d,n}|\beta_{1:K}, z_{d,n} = k) \\
&= \theta_{d,k}\beta_{k,w_{d,n}}
\end{aligned}$$

Next, integrating out the cell type proportions $\theta_d$ and gene expression $\beta_k$ yields an integrand independent of the other assignments and genes. For brevity, we use the shorthand notation $z_{d,n} = k$ as $z_{d,n}$:

$$\begin{aligned}
p(z_{d,n}|z_{-(d,n)}, w) &\propto p(z_{d,n}, w_{d,n}|z_{-(d,n)}, w_{-(d,n)}) \\
&\propto \int_{\beta_k}\int_{\theta_d} p(\theta_d, \beta_k, z_{d,n}, w_{d,n}|z_{-(d,n)}, w_{-(d,n)}) \\
&= \int_{\beta_k}\int_{\theta_d} p(z_{d,n}, w_{d,n}|\theta_d, \beta_k)p(\theta_d|z_{d,-n})p(\beta_k|z_{-(d,n)}, w_{-(d,n)}) \\
&= \int_{\beta_k}\int_{\theta_d} \theta_{d,k}\beta_{k,w_{d,n}} p(\theta_d|z_{d,-n})p(\beta_k|z_{-(d,n)}, w_{-(d,n)}) \\
&= \left(\int_{\theta_d} \theta_{d,k} p(\theta_d|z_{d,-n})\right)\left(\int_{\beta_k} \beta_{k,w_{d,n}} p(\beta_k|z_{-(d,n)}, w_{-(d,n)})\right)
\end{aligned}$$

Each of the above two terms represents the expectation of posterior Dirichlet distributions. Thus, the final algorithm is given as

$$p(z_{d,n} = k|z_{-(d,n)}, \boldsymbol{w}) = \left(\frac{N_{(\cdot)dk}^{-(d,n)} + \alpha}{N_{(\cdot)d(\cdot)} + K\alpha}\right)\left(\frac{N_{i(\cdot)k}^{-(d,n)} + \eta}{N_{(\cdot)(\cdot)k}^{-(d,n)} + V\eta}\right) \tag{3}$$

where,

- $N_{gdk}$: Number of genes of type $g$ in spot $d$ assigned to topic $k$.
- $N_{gdk}^{-(d,n)}$: The count $N_{gdk}$ excluding the contribution of gene $w_{d,n}$.

A new value for $z_{d,n}$ is sampled for each gene $w_{d,n}$ during every iteration of Gibbs sampling. The sampler runs for a burn-in period of 1500 iterations to allow it to reach convergence, after which $\theta_d$ and $\beta_k$ are estimated from $\mathbf{z}$ as follows:

$$\theta_{d,k} = \frac{N_{(\cdot)dk} + \alpha}{N_{(\cdot)d(\cdot)} + K\alpha}$$

$$\beta_{k,g} = \frac{N_{g(\cdot)k} + \eta}{N_{(\cdot)(\cdot)k} + V\eta}$$

### 2 Deconvolution by incorporating gene network structures with gnSPADE

The algorithm described above assumes that genes function independently. However, it is well known that genes do not act in isolation; rather, they operate within complex networks to carry out their biological functions. As a result, gene expression is often correlated. Incorporating this network-based correlation structure has the potential to enhance the accuracy of deconvolution results. To do so, gnSPADE defines a Markov random field (MRF)[5, 6] over the latent cell type layer. Given a spot $d$ with $N_d$ UMI counts, we examine all transcript pairs $(w_{d,i}, w_{d,j})$. If the two transcripts from different genes are known to be correlated based on prior knowledge (e.g., co-expression or functional interaction), we introduce an undirected edge between their respective latent cell type assignments $(z_{d,i}, z_{d,j})$. This results in an undirected graph $\mathcal{G}_d$, where the nodes correspond to the latent variables $\mathbf{z}_d = \{z_{d,n}\}_{n=1}^{N_d}$, and edges connect pairs of assignments associated with correlated genes.

To convert the graph $\mathcal{G}_d$ into an MRF, we define binary edge potentials that favor similar assignments for correlated genes. Specifically, the potential between a connected pair $(i, j) \in \mathcal{P}_d$ is given by:

$$\phi(z_{d,i}, z_{d,j}) = \exp\left\{\frac{\sum_{(i,j)\in\mathcal{P}} \mathbb{I}(z_{d,i} = z_{d,j})}{|\mathcal{P}_d|}\right\} \tag{4}$$

Here, $\mathcal{P}_d$ denotes the set of all edges in the gene correlation graph $\mathcal{G}_d$ for spot $d$, and $|\mathcal{P}_d|$ indicates the total number of such edges. The function $\mathbb{I}(\cdot)$ is the indicator function, which encourages correlated genes to be assigned to the same latent cell type. Under this formulation, the joint probability of the latent cell type assignments within spot $d$ becomes:

$$\begin{aligned} p(\mathbf{z}_d|\boldsymbol{\theta}, \lambda) &= \prod_{i=1}^{N_d} p(z_{d,i}|\boldsymbol{\theta}) \prod_{(i,j)\in\mathcal{P}} \phi(z_{d,i}, z_{d,j})^{\lambda} \\ &= \prod_{i=1}^{N_d} p(z_{d,i}|\boldsymbol{\theta}) \exp\left\{\lambda \frac{\sum_{(i,j)\in\mathcal{P}} \mathbb{I}(z_{d,i} = z_{d,j})}{|\mathcal{P}_d|}\right\} \end{aligned} \tag{5}$$

where $\lambda \geq 0$ is a regularization parameter that controls the strength of the MRF prior. Then we have:

$$p(z_{d,n} = k|z_{-(d,n)}, \boldsymbol{w}) \propto \left(\frac{N_{(\cdot)dk}^{-(d,n)} + \alpha}{N_{(\cdot)d(\cdot)} + K\alpha}\right)\left(\frac{N_{i(\cdot)k}^{-(d,n)} + \eta}{N_{(\cdot)(\cdot)k}^{-(d,n)} + V\eta}\right) \exp\left\{\lambda \frac{\sum_{j\in\mathcal{N}_{di}} \mathbb{I}(z_{d,j} = k)}{|\mathcal{N}_{di}|}\right\} \tag{6}$$

where $\mathcal{N}_{di}$ denotes the genes that are labeled to be similar to gene $i$ in the $d$th spot, $|\mathcal{N}_{di}|$ is the number of transcript counts in $\mathcal{N}_{di}$. When $\lambda = 0$, gnSPADE reduces to the standard LDA model in equation (3), in which each $z_{d,n}$ is conditionally independent given $\theta_d$. In contrast, gnSPADE explicitly encourages the assignments of correlated genes to align, introducing structured dependencies that reflect biological priors.

### 3 Model based data simulation

To generate pseudo cell type expression profiles for 100 genes (gene1, gene2, ..., gene100) for each $\beta_k$ given $k = 1, 2, 3, 4$, we first set 4 groups correlated genes, using 'mvrnorm()' function in R with corvariance matrix to generate correlated dirichlet prior $\boldsymbol{\alpha}$ used in 'gtools::rdirichlet()'. To simulate cell type proportions $\boldsymbol{\theta}$, we applied 'gtools::rdirichlet(n_spots, rep(1/K, K))' to generate cell type proportions across all spots. Assuming an average total count of 1000 per spot, we generated total counts using a Poisson distribution. For each spot, we first sampled the number of counts contributed by each cell type from a multinomial distribution parameterized by $\theta_d$, and then sampled gene counts for each cell type from a multinomial distribution based on the corresponding $\beta_k$.

### 4 Gene network used in the model-free data simulation

#### 4.1 For the MPOA data

We input all 135 genes into STRING[7], an algorithm to extract the protein-protein interaction (PPI) network, with the minimum required interaction score 0.4, and interaction sources: Textmining,

Databases, Co-expression, Neighborhood, Gene Fusion and Co-occurrence , to get the gene network.

##### 4.2 For the MK data

We input all 307 genes into STRING[7], with the minimum required interaction score 0.9, and interaction sources: Textmining, Databases, Co-expression, Neighborhood, Gene Fusion and Co-occurrence, to get the gene network.

#### 5 Real data pre-processing

##### 5.1 MOB ST data

We obtained mouse olfactory bulb (MOB) datasets from the original publication, focusing on MOB replicate #8. Coarse clustering annotations were obtained from STdeconvolve. To ensure a direct comparison with STdeconvolve, We followed the same preprocessing steps. First, we removed genes with fewer than 100 reads detected across spots, and excluded spots with fewer than 100 total gene counts. This filtering process resulted in a cleaned dataset containing 260 spots and 7,365 genes. Next, we selected 255 overdispersed genes using the default generalized additive model (basis = 5) and applied multiple testing with an adjusted p-value $< 0.05$. We input selected 255 genes into STRING[7], with the minimum required interaction score 0.7, and interaction sources: Textmining, Experiments, Databases, Co-expression, Neighborhood, Gene Fusion and Co-occurrence. We set gene connectivity by considering SCC $> 0.5$ in the gene network. Then, we fit the model using integer values of $K$ from 2 to 18 and selected $K = 7$, which minimized perplexity and resulted in fewer rare cell types with mean spot proportions below 5%.

##### 5.2 10x Visium data

A 10x Visium dataset of a coronal section of the mouse cortex was obtained. We removed spots with fewer than 100 gene counts and genes with fewer than 100 total counts, resulting in a dataset with 2,702 pixels and 13,548 genes. To retain a diverse set of biologically significant genes, we only removed genes detected in fewer than 1% or in 100% of spots. From the remaining genes, we selected the top 1,000 most significant overdispersed genes using a generalized additive model (basis = 5) with an adjusted p-value$< 0.05$. We input the top 1,000 genes into STRING[7], with the minimum required interaction score 0.9, and interaction sources: Textmining, Databases, Co-expression, Neighborhood, Gene Fusion and Co-occurrence, to get the gene network. We then fit gnSPADE with integer values of $K$ ranging from 8 to 20 and selected $K = 13$ based on its lower perplexity and smaller number of 'rare' cell types, defined as those with mean spot proportions below 5%.

##### 5.3 DBiT-seq data

We obtained the DBiT-seq dataset of an E11 mouse embryo lower body sample (GSM4364242_E11-1L) from the original publication[8]. After removing spots with fewer than 100 gene counts, genes with fewer than 100 total counts, and mitochondrial genes, we were left with a filtered dataset containing 1,831 spatial locations and 7,171 genes. We then performed feature selection, identifying the top 1,000 most significantly overdispersed genes using a generalized additive model (basis = 5) with an adjusted p-value$< 0.05$. Genes were retained if they were detected in more than 1% but fewer than 100% of the spots. We input the top 1,000 genes into STRING[7], with the minimum required interaction score 0.4, and interaction sources: Co-expression, Neighborhood, and Co-occurrence. We then applied SCC $> 0.5$ to set gene connectivity in the gene network. gnSPADE was fit with integer values of K ranging from 8 to 20, and $K = 13$ was selected based on its lower perplexity and a smaller number of 'rare' cell types (defined as those with mean spot proportions $< 5\%$). This value of $K$ corresponds to the number of transcriptional clusters identified for this sample in the original publication.

##### 5.4 PDAC ST data

We obtained the PDAC dataset from the original publication[9]. For comparison with CARD-free, we used the marker gene lists provided by CARD-free, resulting in 1,379 marker genes that were present in the count matrix. To ensure a fair comparison across different deconvolution methods, we standardized the analysis by implementing deconvolution on the same scaled count matrix with 428 spots and 1,379 genes, setting $K = 20$, corresponding to the number of cell types in the marker gene lists. We input the 1,379 marker genes into STRING[7], with the minimum required interaction score 0.7, and interaction sources: Textmining, Databases, Co-expression, Neighborhood, Gene Fusion and Co-occurrence, to get the gene network.

### 6 Supplementary Figures

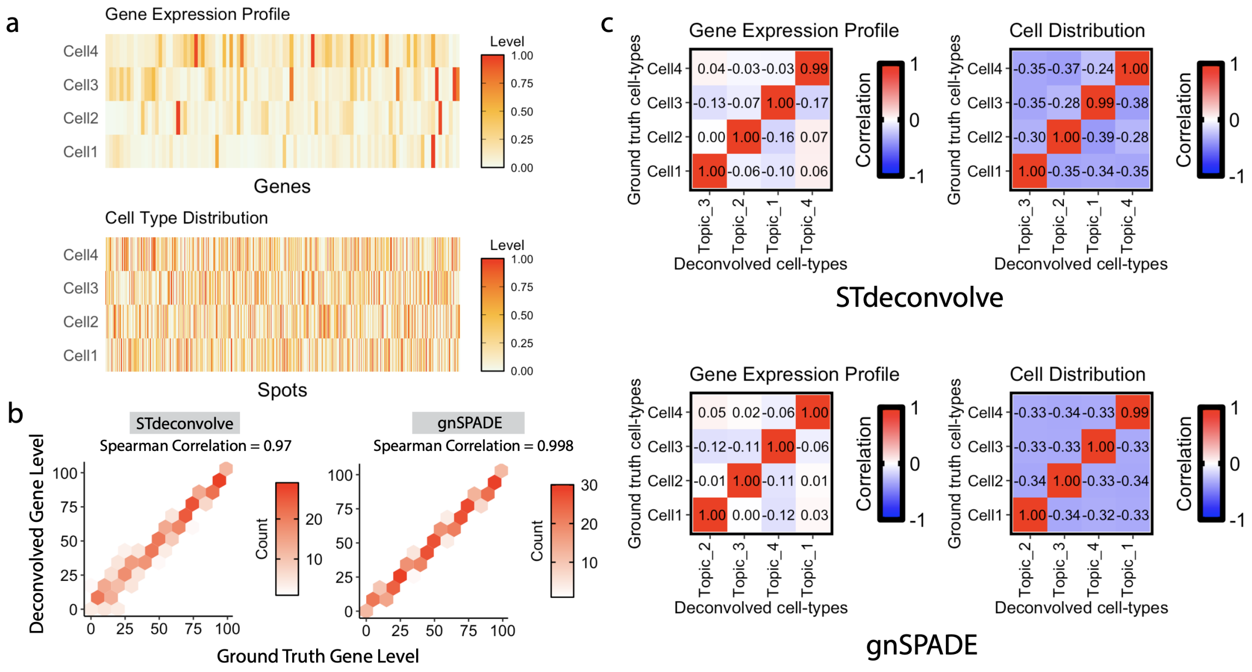


Figure S1: **Comparison between STdeconvolve and gnSPADE in the General gene related synthetic data.** **a** Ground truth gene expression profiles for each cell type (top) and ground truth cell type proportions across all spatial spots (bottom) in the general gene correlation simulation. **b** Gene ranking consistency between ground truth and deconvolved cell-type transcriptional profiles. Each point represents the rank of a gene's expression in the deconvolved profile versus its rank in the corresponding ground truth profile. The associated SCC quantifies ranking agreement. Left: STdeconvolve; Right: gnSPADE. **c** PCC comparisons between ground truth and deconvolved results for the cell–type–specific simulation. Top: STdeconvolve; Bottom: gnSPADE. Left: Transcriptional profiles (ground truth vs. deconvolved); Right: Cell type compositions (ground truth vs. deconvolved).

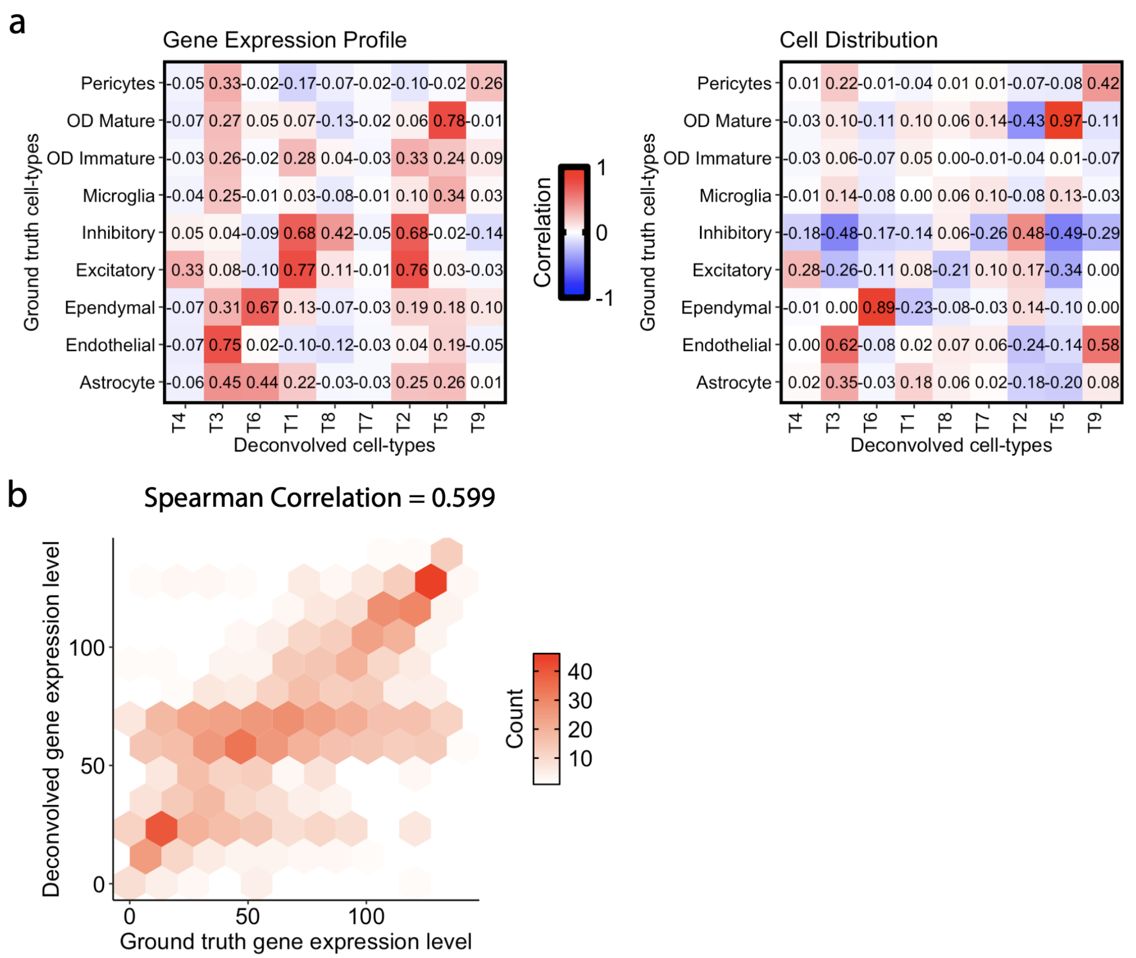


Figure S2: **SpiceMix deconvolution results for MPOA. a** Heatmaps of Pearson correlation coefficients comparing ground truth with deconvolved results. **b** Gene expression ranking consistency. Each gene is ranked by expression level in the deconvolved transcriptional profiles and compared to its rank in the matched ground truth profile. The corresponding SCC = 0.599 quantifies rank similarity.

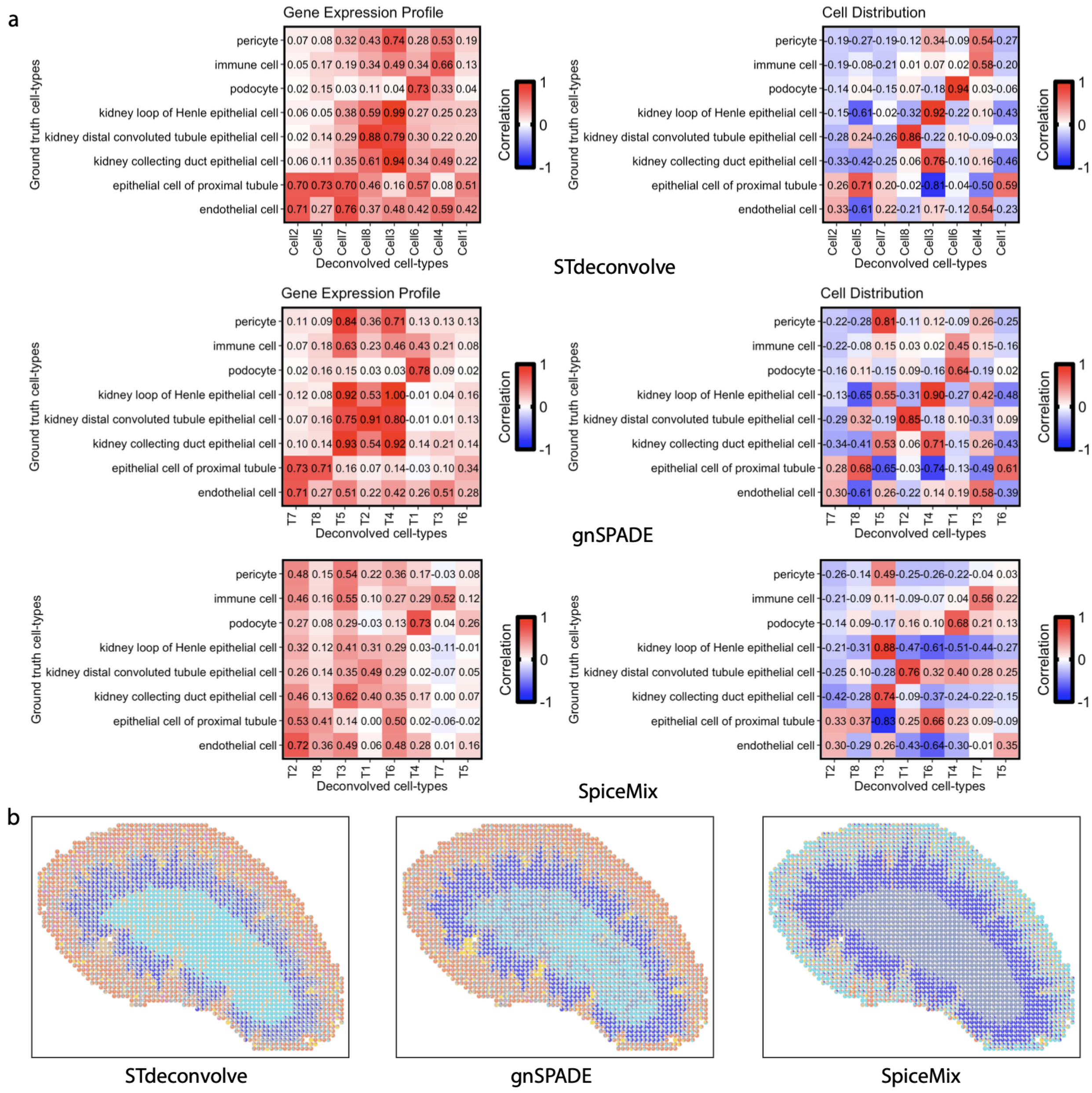


Figure S3: **Comparison between STdeconvolve, gnSPADE and SpiceMix in the MK data. a** (Left) Pearson's correlation between the transcriptional profiles of the 8 ground truth cell types in the MERFISH MK data and the corresponding 8 deconvolved cell types. (Right) Pearson's correlation between the simulated grid proportions of the 8 ground truth cell types and the 8 deconvolved cell types. (Top: STdeconvolve; Middle: gnSPADE; Bottom: SpiceMix). **b** Predicted grid proportions of 8 deconvolved cell types from STdeconvolve, gnSPADE and SpiceMix.

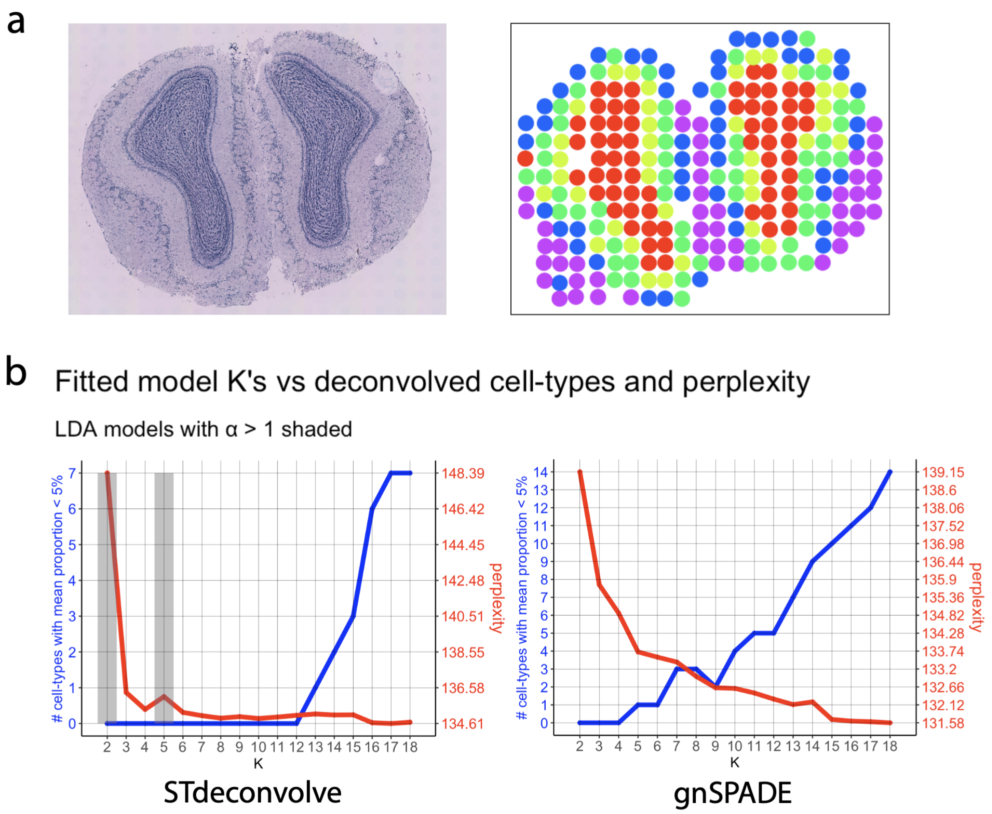


Figure S4: **Additional plots related to the MOB data.** **a** (Left) H&E-stained image of the mouse olfactory bulb (MOB) tissue section. (Right) Visualization of the corresponding MOB spots, colored by their transcriptional cluster memberships mapped to spatial locations. **b** Plot of perplexity and the # of cell types with mean proportional $<5\%$ under different numbers of cell types $K$ for STdeconvolve and gnSPADE.

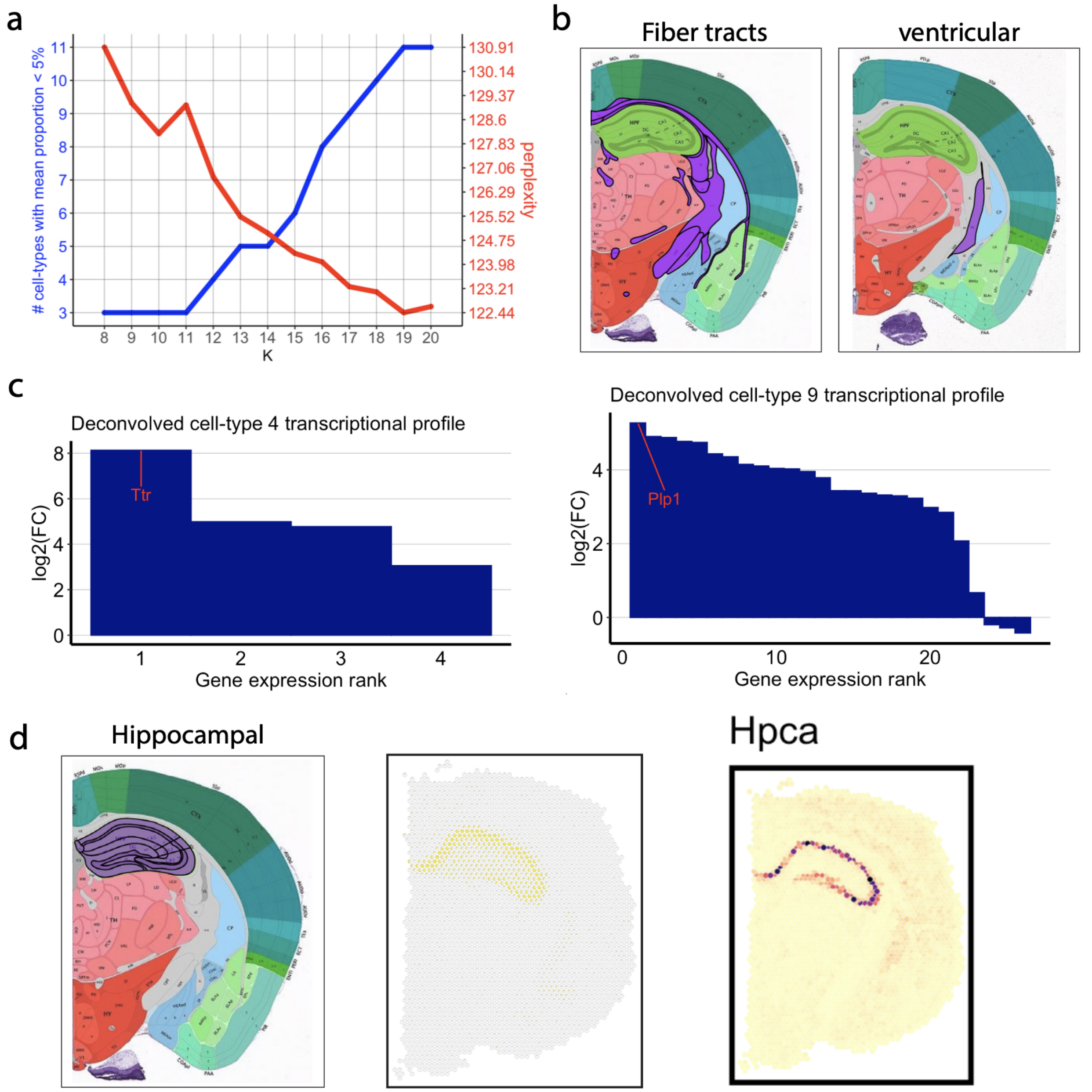


Figure S5: **Additional plots related to the 10× Visium mouse brain data. a** Plot of perplexity and the # of cell types with mean proportional < 5% under different numbers of cell types $K$ for gnSPADE. **b** Histological annotation regions. (Left: fiber tracts; Right: ventricular). **c** Log2 fold-change analysis of deconvolved gene transcriptional profiles for each deconvolved cell type compared to the mean deconvolved expression of the other 13 cell types (Left: for cell type 4; Right: for cell type 9.), matched with histological annotation regions. **d** From left to right: histological annotation regions, highlighted deconvolved cell types corresponding to the annotation regions, and the corresponding top differentially expressed gene counts across all spots (in Hippocampus).

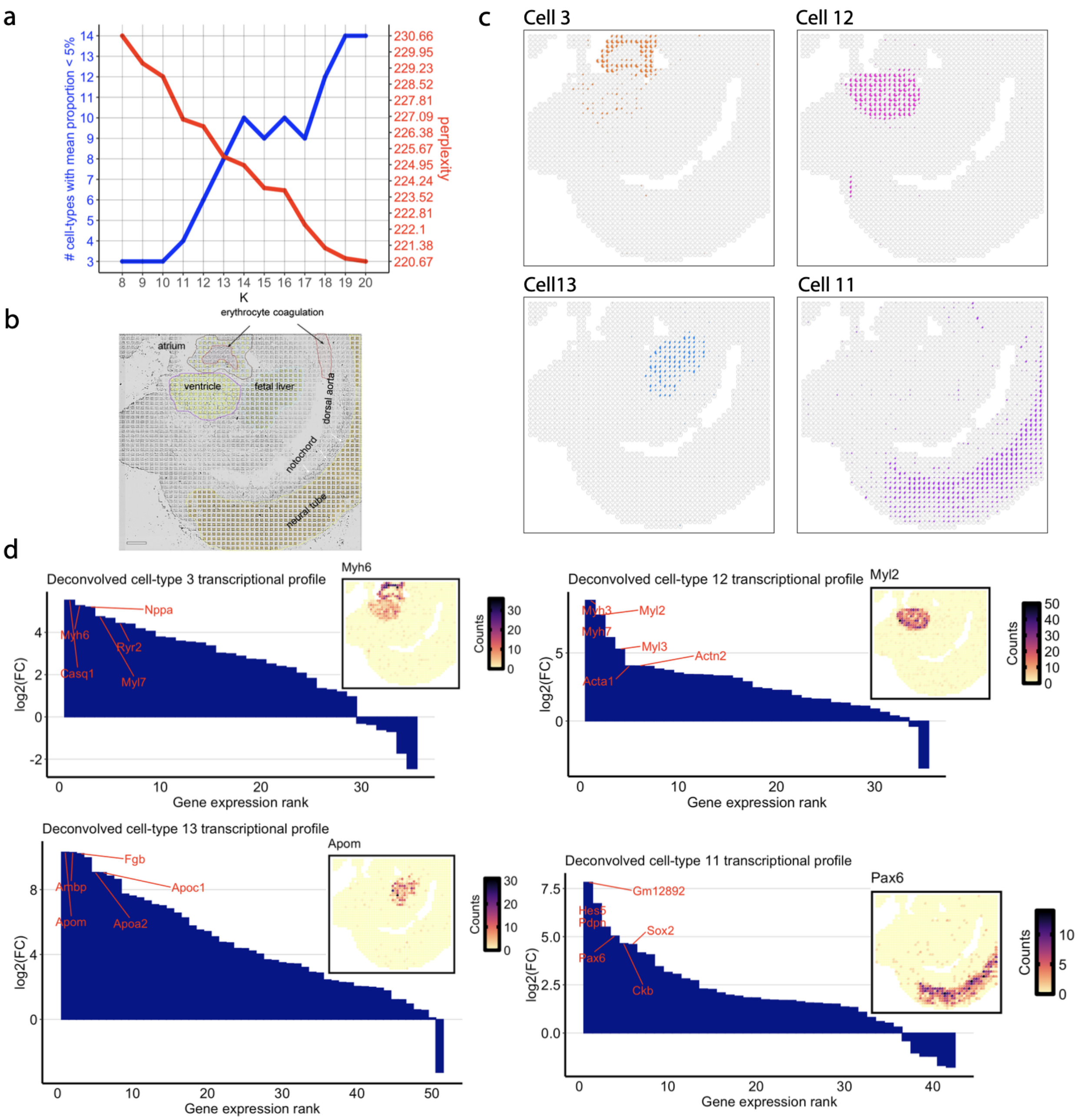


Figure S6: **Spots proportions and log2 fold of transcriptional profiles of the selected deconvolved cell-types in DBiT-seq data of the E11 mouse embryo lower tail section.** **a** The relationship between the number of cell types $K$ and the number of "rare" cell types, along with perplexity scores for gnSPADE. **b** Tissue types identified from the original paper's clustering[8]. **c** Visualization of the spot proportions for select deconvolved cell types, Cell1 (atrium), Cell12 (ventricle), Cell13 (fetal liver), and Cell5 (neural tube), corresponding to annotations from a previous publication[8]. **d** Log2 fold-change analysis of the deconvolved transcriptional profile of these cell types with respect to the mean expression of the other 12 deconvolved cell types. The expression of the top differentially expressed genes in each deconvolved transcriptional profile is visualized in the original tissue (inset).

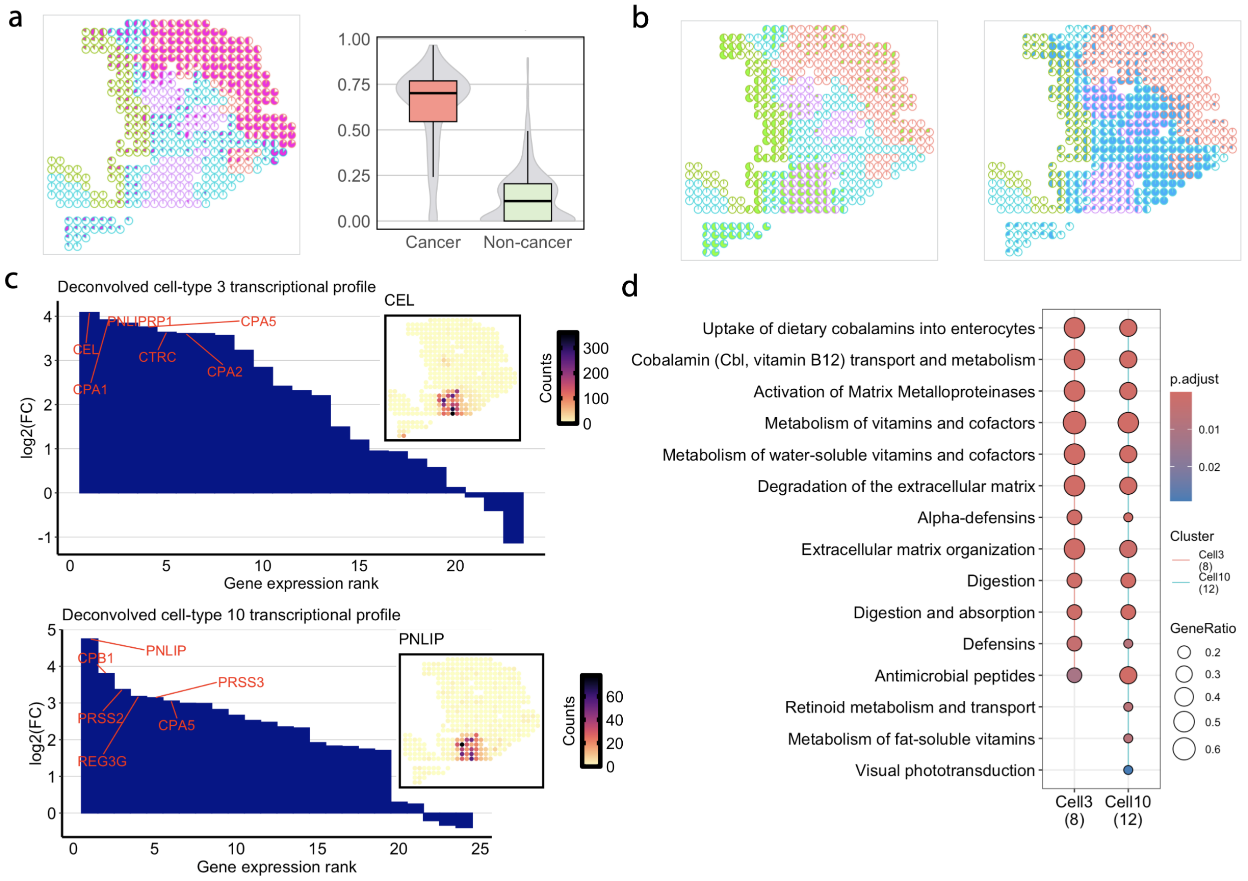


Figure S7: **SpiceMix deconvolution (a-b) and analysis of acinar cells in the PDAC ST data with gnSPADE (c-d).** **a** (Left) Highlights of the identified deconvolved cell types for tumors from SpiceMix. (Right) Comparison of cell-type proportions inferred by SpiceMix in cancer regions versus non-cancer regions. **b** Visualization of major deconvolved cell types in different regions. (Left) Ductal region. (Right) Stroma region. **c** Log2 fold-change of Cell3 and Cell10 (Acinar similar cell) transcriptional profiles relative to the average expression of the other 19 deconvolved cell types, with visualization of the top differentially expressed genes in each deconvolved cell type. (Top: Cell3; Bottom: Cell10) **d** Pathway enrichment analysis of top expressed genes from Cell3 and Cell10 based on the Reactome pathway. The number below each cell type represents the number of overlapped genes between the top expressed genes of that cell type and all genes in the Reactome collection. The GeneRatio represents the ratio of the overlap size between top expressed genes of a specific cell type and a particular pathway gene set to the total overlap with all Reactome pathway members.

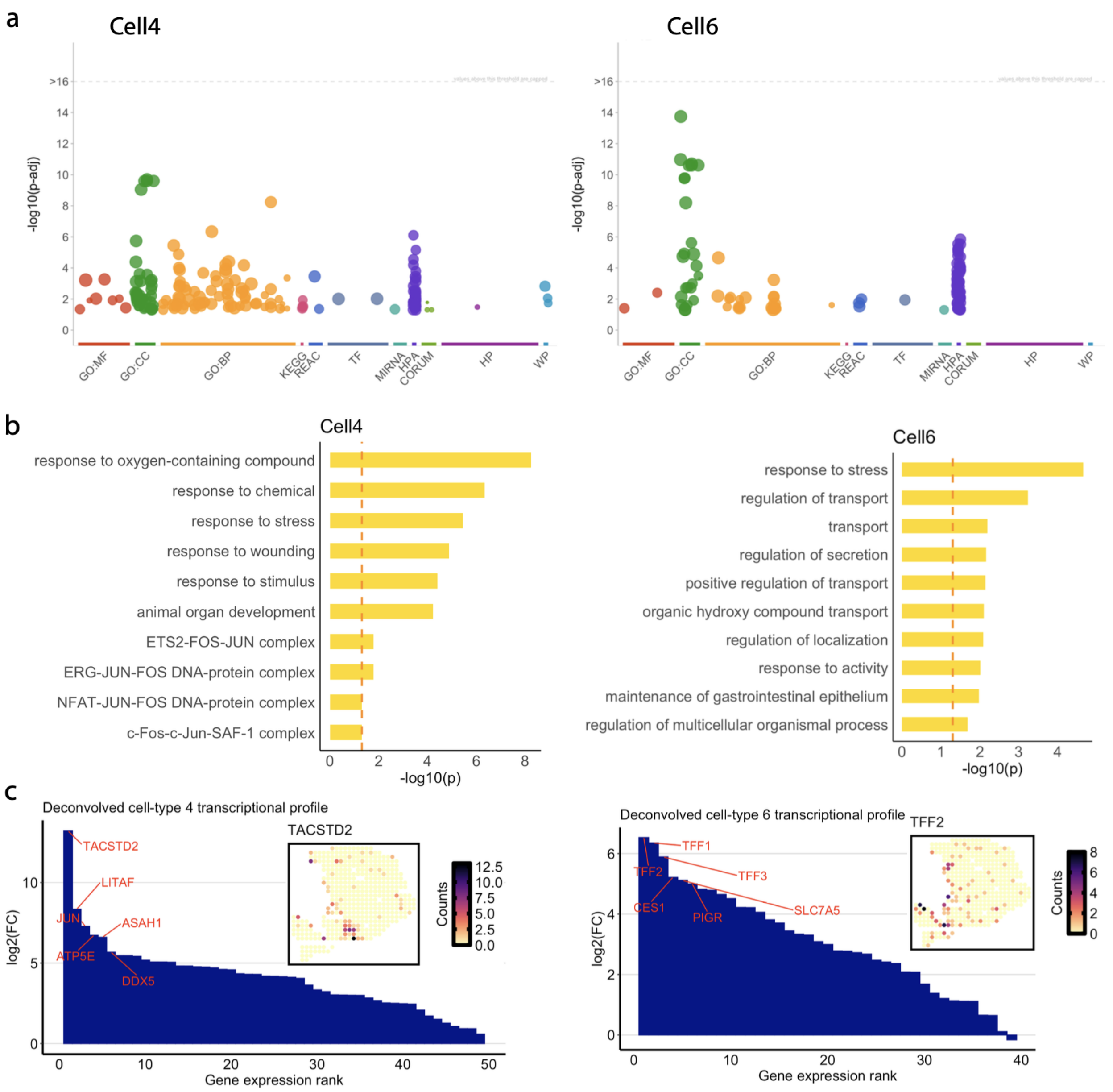


Figure S8: **Analysis of Acinar cells in the deconvolution of the PDAC ST data.** **a** Visualization of adjusted p-values for pathway enrichment analysis cross GO:BP, GO:MF, GO:CC, KEGG, REAC, TF, MIRNA, CORUM, HP, HPA, and WP pathways [10]. **b** Visualization of the top 10 pathways obtained from **a**. **c** Log2 fold-change of Cell4 and Cell6 (Acinar similar cell) transcriptional profiles relative to the average expression of the other 19 deconvolved cell types, with visualization of the top differentially expressed genes in each deconvolved cell type. (Left: Cell4; Right: Cell6)

### References


[1] D. M. Blei, A. Y. Ng, and M. I. Jordan, "Latent dirichlet allocation," *Journal of Machine Learning Research*, vol. 3, no. Jan, pp. 993–1022, 2003.

[2] B. F. Miller, F. Huang, L. Atta, A. Sahoo, and J. Fan, "Reference-free cell type deconvolution of multi-cellular pixel-resolution spatially resolved transcriptomics data," *Nature Communications*, vol. 13, no. 1, p. 2339, 2022.

[3] T. L. Griffiths and M. Steyvers, "Finding scientific topics," *Proceedings of the National Academy of Sciences*, vol. 101, no. suppl_1, pp. 5228–5235, 2004.

[4] D. M. Blei, "Mixed-membership models (and an introduction to variational inference)," *Course notes for Foundations of Graphical Models. Nov*, 2015.

[5] P. Xie, D. Yang, and E. Xing, "Incorporating word correlation knowledge into topic modeling," in *Proceedings of the 2015 conference of the north American chapter of the association for computational linguistics: human language technologies*, pp. 725–734, 2015.

[6] J. Qiang, P. Chen, T. Wang, and X. Wu, "Topic modeling over short texts by incorporating word embeddings," in *Advances in Knowledge Discovery and Data Mining: 21st Pacific-Asia Conference, PAKDD 2017, Jeju, South Korea, May 23-26, 2017, Proceedings, Part II 21*, pp. 363–374, Springer, 2017.

[7] D. Szklarczyk, A. L. Gable, K. C. Nastou, D. Lyon, R. Kirsch, S. Pyysalo, N. T. Doncheva, M. Legeay, T. Fang, P. Bork, *et al.*, "The string database in 2021: customizable protein–protein networks, and functional characterization of user-uploaded gene/measurement sets," *Nucleic Acids Research*, vol. 49, no. D1, pp. D605–D612, 2021.

[8] Y. Liu, M. Yang, Y. Deng, G. Su, A. Enninful, C. C. Guo, T. Tebaldi, D. Zhang, D. Kim, Z. Bai, *et al.*, "High-spatial-resolution multi-omics sequencing via deterministic barcoding in tissue," *Cell*, vol. 183, no. 6, pp. 1665–1681, 2020.

[9] R. Moncada, D. Barkley, F. Wagner, M. Chiodin, J. C. Devlin, M. Baron, C. H. Hajdu, D. M. Simeone, and I. Yanai, "Integrating microarray-based spatial transcriptomics and single-cell rna-seq reveals tissue architecture in pancreatic ductal adenocarcinomas," *Nature Biotechnology*, vol. 38, no. 3, pp. 333–342, 2020.

[10] L. Kolberg, U. Raudvere, I. Kuzmin, J. Vilo, and H. Peterson, "gprofiler2–an r package for gene list functional enrichment analysis and namespace conversion toolset g: Profiler," *F1000Research*, vol. 9, pp. ELIXIR–709, 2020.
